# High-content analysis and Kinetic Image Cytometry identify toxic and epigenotoxic effects of HIV antiretrovirals on human iPSC-neurons and primary neural precursor cells

**DOI:** 10.1101/2020.09.05.284422

**Authors:** Alyson S. Smith, Soneela Ankam, Chen Farhy, Lorenzo Fiengo, Ranor C.B. Basa, Kara L. Gordon, Charles T. Martin, Alexey V. Terskikh, Kelly L. Jordan-Sciutto, Jeffrey H. Price, Patrick M. McDonough

## Abstract

Despite viral suppression due to combination antiretroviral therapy (cART), HIV-associated neurocognitive disorders (HAND) continue to affect half of people with HIV, suggesting that certain antiretrovirals (ARVs) may contribute to HAND. We examined the effects of nucleoside/nucleotide reverse transcriptase inhibitors tenofovir disproxil fumarate (TDF) and emtricitabine (FTC) and the integrase inhibitors dolutegravir (DTG) and elvitegravir (EVG) on viability, structure, and function of glutamatergic neurons (a subtype of CNS neuron involved in cognition) derived from human induced pluripotent stem cells (hiPSC-neurons), and primary human neural precursor cells (hNPCs), which are responsible for neurogenesis. Using automated digital microscopy and image analysis (high content analysis, HCA), we found that DTG, EVG, and TDF decreased hiPSC-neuron viability, neurites, and synapses after seven days of treatment. Analysis of hiPSC-neuron calcium activity using Kinetic Image Cytometry (KIC) demonstrated that DTG and EVG also decreased the frequency and magnitude of intracellular calcium transients. Longer ARV exposures and simultaneous exposure to multiple ARVs increased the magnitude of these neurotoxic effects. Using the Microscopic Imaging of Epigenetic Landscapes (MIEL) assay, we found that TDF decreased hNPC viability and changed the distribution of histone modifications that regulate chromatin packing, suggesting that TDF may reduce neuroprogenitor pools important for CNS development and maintenance of cognition in adults. This study establishes human preclinical assays that can screen potential ARVs for CNS toxicity to develop safer cART regimens and HAND therapeutics.

## Introduction

About 40 million people live with human immunodeficiency virus (HIV) infection globally, and there are ~1.7 million new infections per year^1^. Combination antiretroviral therapy (cART), which involves simultaneous treatment with 2-4 antiretrovirals (ARVs), suppresses HIV replication, prevents progression to acquired immunodeficiency syndrome (AIDS), and extends life expectancy in adults with HIV to near normal^2,3^. Preexposure prophylaxis with cART can also prevent transmission of HIV to HIV-negative adolescents and adults at risk for behaviorally acquiring HIV^4–8^, as well as mother-to-child transmission^9,10^. cART has greatly reduced the incidence of AIDS-defining illnesses that affect the central nervous system (CNS), including opportunistic infections, primary CNS lymphoma, and HIV-associated dementia (HAD)^11–14^. However, the post-cART era has seen increased incidence of milder forms of HIV-associated neurocognitive disorder (HAND), including mild cognitive disorder (MND) and asymptomatic neurocognitive impairment (ANI)^15,16^. HAND affects approximately 50% of people living with HIV, including patients with undetectable viral loads^17,18^ and can impact quality of life, employment, treatment adherence, and survival^19–21^. HAND can also increase the risk for progression to more severe cognitive impairment^22,23^.

While many factors likely contribute to cognitive impairment in people with HIV on cART, there is growing concern that ARV neurotoxicity contributes^24^. In pigtail macaques, rodents, and cultured rodent neurons, ARV exposure can cause oxidative stress, endoplasmic reticulum stress, mitochondrial dysfunction, loss of neurites and synapses, and/or neuronal cell death^25–28^. In people with HIV, cART can reduce neuronal metabolite levels^29^, the volume and structural integrity of cortical white matter^30,31^, and cognitive reserve as measured by functional MRI^32^. While some studies link cART regimens with high CNS penetration effectiveness (CPE) with improved cognition^33^ or no effect on cognition^34^, others link high CPE with worse cognitive outcomes^35,36^, including a study with 61,938 people with HIV^37^. Treatment interruption in HIV+ individuals with stable immune function can also improve neurocognitive performance for extended time periods^38^. People with HIV must remain on cART for their lifetime to maintain viral suppression^39,40^, leaving them vulnerable to increased neurotoxicity during neurodevelopment and aging. Indeed, children with HIV receiving cART have reduced cognition^41–43^, and HIV-children who received perinatal cART experience developmental delays^44^. cART also increases production and reduces clearance of Alzheimer’s disease-associated beta amyloid peptides in vitro^45–47^ and in patients^48,49^, which may accelerate CNS aging and cognitive decline.

In previous research using HCA and KIC, we found that tamoxifen, an anti-cancer agent linked to post-chemotherapy cognitive impairment, reduces synapses and calcium transient activity of primary rat hippocampal neurons, establishing that these in vitro techniques can test agents for potential negative impacts on cognition^50^. In this study, we aimed to develop human preclinical neurosafety testing platforms to screen ARVs for a broad range of potential neurotoxic and neurodevelopmental effects. We first developed HCA and KIC assays using glutamatergic neurons differentiated from human induced pluripotent stem cells (hiPSC-neurons), which provide an in vitro model system with similar gene expression, cell biology, and electrophysiology to neurons in the human brain in which to test directly for compound toxicity^51–53^. We also developed an assay to test ARVs for potential effects on neurogenesis, a process by which human neural precursor cells (hNPCs) differentiate to neurons to support central nervous system development and cognitive function in adults^54,55^. For this, we used the MIEL assay, a multiparametric approach to identify changes in histone acetylation and methylation patterns using HCA of immunofluorescence images^56^.

Using HCA and KIC methods on hiPSC-neurons, we found that the nucleoside/nucleotide reverse transcriptase inhibitors tenofovir disproxil fumarate (TDF) and emtricitabine (FTC) and the integrase inhibitors dolutegravir (DTG) and elvitegravir (EVG) altered key neuronal structures and functions to varying extents, and that combinations of these ARVs had additive effects. Additionally, the MIEL assay showed that TDF reduced viability and altered histone acetylation and methylation in hNPCs, indicating an epigenotoxic effect. Our findings suggest novel preclinical research strategies to test candidate ARVs in vitro and to aid in identifying ARVs with reduced neurocognitive effects.

## Materials and Methods

### Cell Culture

For hiPSC-neuron experiments, imaging-quality polystyrene 384-well plates (Greiner Bio-One #781090; Frickenhausen, Germany) were coated with 0.1% polyethyleneimine and 0.028 mg/mL growth factor-reduced Matrigel (Corning Life Sciences #354230, Tewksbury, MA, USA). hiPSC-derived glutamatergic neurons (iCell GlutaNeurons; Fujifilm CDI #C1060, Madison, WI, USA) were seeded at a density of 60,000 live cells/cm^2^ and were maintained per the manufacturer’s instructions for a total of 14 days before assay.

For hNPC experiments, imaging-quality polystyrene 384-well plates (Greiner Bio-One #781090; Frickenhausen, Germany) were coated with 0.028 mg/mL growth factor-reduced Matrigel (Corning Life Sciences #354230, Tewksbury, MA, USA). Fetal hNPCs (Thermo Fisher Scientific # A15654; Waltham, MA, USA) were seeded at a density of 18,200 live cells/cm^2^ and were maintained in differentiation medium per the manufacturer’s instructions for a total of 7 days before assay.

### Test Compounds

hiPSC-neuron cultures were treated with ARVs or ARV combinations for 1 or 7 days prior to assays. The following ARVs were used alone or in combination in this study: dolutegravir (DTG; Toronto Research Chemicals #D528800; Toronto, ON, Canada), elvitegravir (EVG; Toronto Research Chemicals #E509000), tenofovir disproxil fumarate (TDF; Toronto Research Chemicals #T018505), emtricitabine (FTC; Toronto Research Chemicals #E525000), DTG+TDF+FTC, EVG+TDF+FTC, and TDF+FTC. Vehicle (0.2% DMSO) and antagonist control treatments (25 μM *para*-nitroblebbistatin; Optopharma Ltd. #DR-N-111; Budapest, Hungary) were also applied to the cells. hNPCs were treated with 0.2% DMSO, ARVs, SAHA (Cayman Chemical #10009929; Ann Arbor, MI, USA), GSK343 (Cayman Chemical #14094), tofacitinib (TOF, Cayman Chemical #11598), or (+)-JQ1 (Cayman Chemical #11187).

### Automated Microscopy

The fixed-endpoint synapse and neurite length assay and microscopic imaging of the epigenetic landscape (MIEL) assay, as well as the live-cell calcium KIC assay were imaged using the IC200-KIC automated microscope (Vala Sciences Inc.; San Diego, CA, USA; Kinetic Image Cytometry® and KIC® are registered trademarks of Vala Sciences Inc.) outfitted with a Plan Apo 20X/0.75 NA objective lens (Nikon Instruments Inc.; Melville, NY, USA). For live-cell assays, the environmental chamber of the IC200-KIC was set to 37°C/5% CO_2_.

### Synapse Density and Neurite Length Assay

After treatment, cells were fixed in 2% paraformaldehyde/1.67% sucrose in HBSS without Ca^++^/Mg^++^, permeabilized in 0.3% Triton X-100 in PBS with Ca^++^/Mg^++^, and blocked in 5% normal goat serum/1% BSA/0.1% Triton X-100. The following primary antibodies were diluted in blocking buffer and applied to cells overnight at 4°C: chicken anti-βIII tubulin (Tuj-1, 1:200; Abcam #ab41489; Boston, MA, USA), rabbit anti-PSD95 (1:200; Thermo Fisher Scientific #51-6900; Waltham, MA, USA), and mouse anti-SV2 (1:150; Developmental Studies Hybridoma Bank; Iowa City, IA, USA). The next day, the following secondary antibody cocktail with Hoechst nuclear stain was made in 2% BSA and applied to the cells for 1 hour at room temperature in the dark: goat anti-chicken IgY Alexa Fluor 555 (1:500; Thermo Fisher Scientific #A21437), goat anti-rabbit IgG Alexa Fluor 647 (1:500; Thermo Fisher Scientific #A21245), goat anti-mouse IgG Alexa Fluor 488 (1:500; Thermo Fisher Scientific #A11029), and 10 μg/mL Hoechst 33342 (Thermo Fisher Scientific #H3570). For each well, a 3-by-3 matrix of images was acquired in each optical channel.

### Calcium KIC Assay

Neurons were incubated in a calcium indicator dye solution consisting of 5 μM Rhod-4 AM (AAT Bioquest #21122; Sunnyvale, CA, USA), 1X PowerLoad (Thermo Fisher Scientific #P10020), 1 μg/mL Hoechst 33342, and 2.5 mM probenecid in phenol red-free BrainPhys (Stemcell Technologies #05791; Vancouver, BC, Canada) for 40 min at 37°C. The neurons were then rinsed with and imaged under phenol red-free BrainPhys. The calcium indicator dye solution and the imaging media did not contain DMSO or ARVs. One field of view was imaged per well, including a single image in the nuclear channel followed by a calcium movie acquired at 4 frames per second for 2 minutes.

### hNPC Microscopic Imaging of Epigenetic Landscape (MIEL) Assay

After treatment, cells were fixed in 2% paraformaldehyde/1.67% sucrose in HBSS without Ca^++^/Mg^++^ and blocked/permeabilized in PBS with Ca^++^/Mg^++^ and 2% BSA/0.5% Triton X-100. The following primary antibodies were diluted in blocking buffer and applied to cells overnight at 4°C to label key histone modifications associated with epigenetic regulation: rabbit H3K9me3 (1:500; Active Motif, 39765; Carlsbad, CA, USA); mouse H3K4me1 (1:100; Active Motif, 39635; Carlsbad, CA, USA); mouse H3K27me3 (1:250; Active Motif, 61017; Carlsbad, CA, USA); and rabbit H3K27ac (1:500; Active Motif, 39133; Carlsbad, CA, USA). Two sets of cells were immunolabeled for the biomarkers: one set with H3K27me3 and H3K27ac and a second set with H3K9me3 and H3K4me1. The next day, the following secondary antibody cocktail with Hoechst nuclear stain was made in 2% BSA and applied to the cells for 1 hour at room temperature in the dark: goat anti-rabbit IgG Alexa Fluor 647 (1:500; Thermo Fisher Scientific #A21245), goat anti-mouse IgG Alexa Fluor 488 (1:500; Thermo Fisher Scientific #A11029), and 10 μg/mL Hoechst 33342 (Thermo Fisher Scientific #H3570).

### Automated Image Analysis

For hiPSC-neuron synapse density, neurite length, and calcium KIC assays, images were analyzed using custom algorithms in CyteSeer software (Vala Sciences Inc.; CyteSeer® is a registered trademark of Vala Sciences Inc.). The NO2 v7.2.4 Neurite Morphology and Synapse Density with Somas algorithm was used to analyze the synapse density and neurite length images. This algorithm identified live neuronal nuclei as relatively large nuclei with diffuse Hoechst staining (as opposed to small, bright, dead nuclei) that colocalize with neuron-specific βIII-tubulin (Tuj-1) staining. The algorithm then identified Tuj-1+ neurites, which includes axons and dendrites. Small neurite fragments and Tuj-1+ cellular debris were excluded, as well as the Tuj-1 positive neuronal cell bodies (somas). To identify synapses, the algorithm first identified SV2+ (presynaptic) and PSD95+ (postsynaptic) puncta with areas between 0.32 and 1.27 μm^2^ (between 3 and 15 pixels^2^ with a pixel size of 0.325 μm). To be considered synapses, the SV2+ puncta centroids were required to be within 1.95 μm (6 pixels) of the nearest neurite and within 0.975 μm (3 pixels) of the nearest PSD95+ puncta centroid. Neuronal viability, neurite, and synapse data are presented as one data point per well, with each data point representing the average value across each of the 9 fields of view acquired per well. Each field of view contained about 200 live neurons in DMSO-treated control wells.

The calcium KIC assays were analyzed with the Neuron Calcium KIC v10.5 algorithm. Neuronal cell bodies were identified as Rhod-4 AM-positive areas associated with live neuron nuclei. The average pixel intensity of Rhod-4 AM signal in each cell body was measured for each of the 480 frames (4 frames per second for two minutes). The resulting functions of Rhod-4 AM average pixel intensity over time were baseline subtracted to determine if each neuron displayed calcium transients during the recording period and to calculate the event frequencies and mean peak amplitudes for each active cell. Event frequency and mean peak amplitude data are presented as one data point per well, with each data point representing the average value of all active neurons in each field of view. Each field of view contained about 200 live neurons in DMSO-treated control wells.

For the MIEL assay, images were analyzed using Acapella 2.6 (PerkinElmer), and image texture features were analyzed as described in Farhy, et al., eLife, 2019^56^. Quadratic discriminant analysis was performed on histone modification texture features using the Excel add-on program Xlstat (Base, v19.06).

### Statistical Analysis

The data were analyzed with Prism (GraphPad Software; San Diego, CA, USA) using ANOVA, followed by either Dunnett’s or Tukey’s post-hoc multiple comparisons test for statistical significance (\**p*<0.05, \*\**p*<0.01, \*\*\**p*<0.001).

## Results

### HCA to characterize HIV ARV effects on neuronal viability and neurites in hiPSC-neurons

To test for ARV neurotoxicity, we exposed hiPSC-neurons to 0.1, 1, or 10 μM of DTG, EVG, TDF, or FTC for seven days. We chose these concentrations to test for neurotoxic effects at levels below, near, and above the plasma Cmax of each ARV (Table 1). We used 25 μM blebbistatin, a nonmuscle myosin II inhibitor^57,58^, as a control compound. We then fixed the hiPSC-neurons and immunolabeled for Tuj-1 (neuron-specific βIII-tubulin), SV2 (presynaptic protein), PSD95 (postsynaptic protein), stained with Hoechst (nuclei), and imaged the cells with the IC200 Image Cytometer to assess neuronal viability and morphology. Images of Tuj-1 and nuclei show significant loss of neurites in hiPSC-neurons treated with 10 μM EVG, indicating a neurotoxic effect, with smaller effects for the other ARVs (Fig. 1A). Image analysis with a CyteSeer algorithm that identifies the nuclei corresponding to live neurons (see Methods) determined that Neuronal Viability (the percentage of live neuronal nuclei) was significantly reduced by 30% at 10 μM DTG, 65% at 10 μM EVG, and 35% at 10 μM TDF (Fig. 1B). We did not observe changes in Neuronal Viability at lower concentrations of TDG, EVG, or TDF, or at any tested concentration of FTC.

**Table 1.** Plasma concentrations of HIV antiretrovirals used in this study.

| **Antiretroviral** | **Abbreviation** | **Drug class** | **Plasma Cmax** |
| --- | --- | --- | --- |
| Dolutegravir | DTG | INSTI | 8 µM |
| Elvitegravir | EVG | INSTI | 3.5 µM |
| Tenofovir disproxil fumarate | TDF | NRTI | 0.6 µM |
| Emtricitabine | FTC | NRTI | 7 µM |

INSTI: integrase strand transfer inhibitor; NRTI: nucleoside/nucleotide reverse transcription inhibitor. Data taken from Food and Drug Administration New Drug Applications for each antiretroviral.

**Figure 1.**
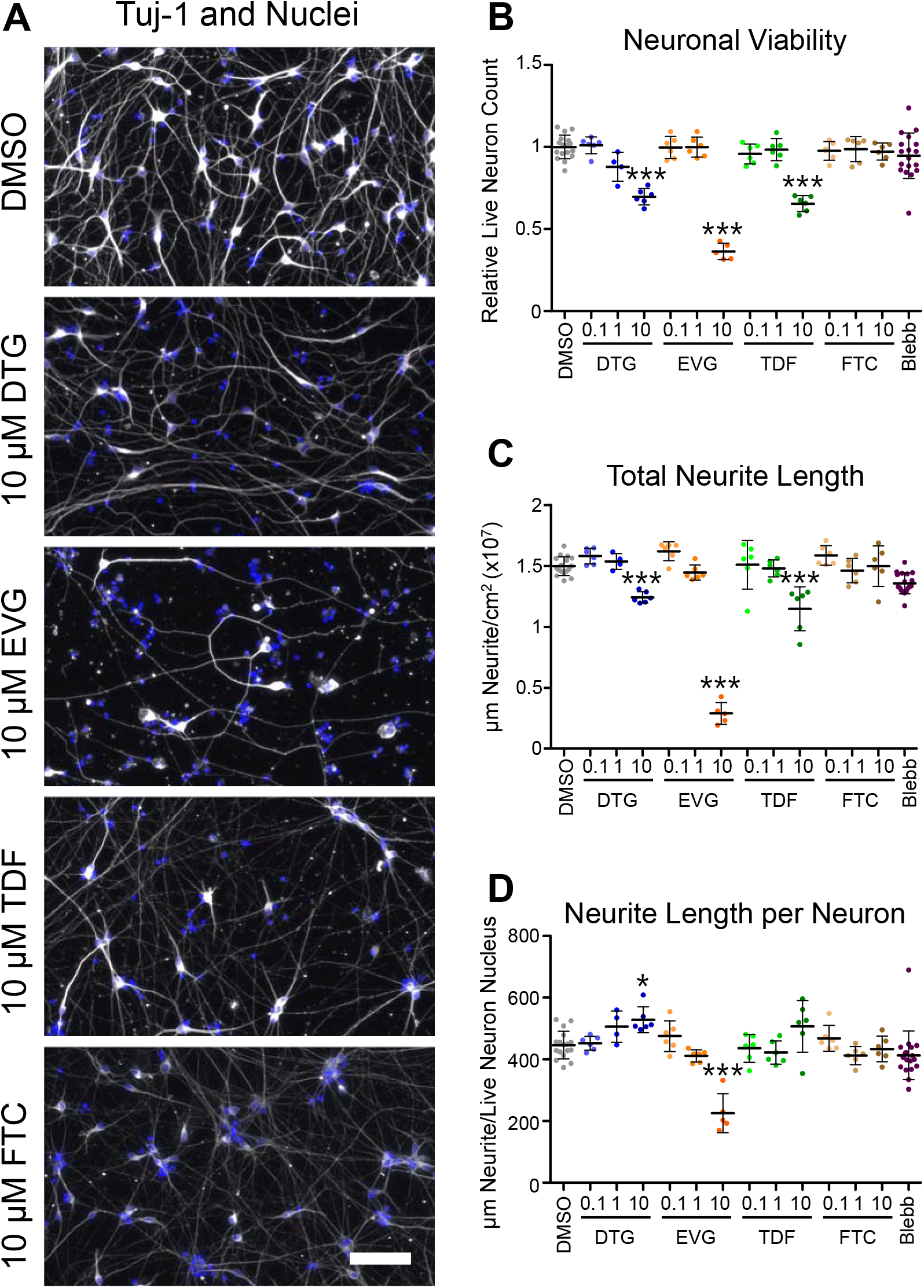
Single ARVs decrease viability and neurite length in hiPSC-neurons in a dose-dependent manner. hiPSC-neurons were exposed to the ARVs for 7 days. (A) Representative images of hiPSC-neurons treated with DMSO alone or 10 μM dolutegravir (DTG), elvitegravir (EVG), tenofovir disproxil fumarate (TDF), or emtricitabine (FTC), fixed, and stained for nuclei (Hoechst, blue) and neuronal somas and neurites (Tuj-1, grayscale). Scale bar = 50 μm. (B-D) Viability and neurite data for hiPSC-neurons treated with DMSO alone, ARVs (0.1, 1, or 10 μM) or blebbistatin (25 μM). (B) Neuronal viability as defined by the number of live neuron nuclei in each well relative to the DMSO control mean. (C) Total neurite length in μm per cm^2^ image area for each condition. (D) Total neurite length per live neuron nucleus for each condition. Each dot represents the average of each measurement from nine images per well. DMSO and blebbistatin: n = 18 wells; ARVs: n = 6 wells. Bars represent mean ± standard deviation. Statistics performed with one-way ANOVA followed by Dunnett’s multiple comparisons test. * p < 0.05, ** p < 0.01, *** p < 0.001.

To further quantify effects of the ARVs, we used a CyteSeer image analysis algorithm that traces Tuj-1-positive neurites and calculates their length. The total neurite length per cm^2^ image area (Total Neurite Length) was significantly reduced by 20% at 10 μM DTG, 80% at 10μM EVG, and 20% at 10μM TDF (Fig. 1C). The Neurite Length per Neuron (Total Neurite Length divided by the number of live neuronal nuclei) was slightly but significantly increased by 20% (p < 0.05) after treatment with 10 μM DTG but was 50% lower after treatment with 10 μM EVG (Fig. 1D). The Total Neurite Length and Neurite Length per Neuron remained similar to DMSO alone at lower concentrations of DTG, EVG, and TDF, and at all tested concentrations of FTC.

Most HIV infections are treated with combination antiretroviral therapy (cART) consisting of two, three, or four ARVs from two or more different classes^2–4^. To test whether cART causes increased neurotoxicity and neurite loss compared to single ARVs, we compared Neuronal Viability and neurite length measurements in hiPSC-neurons treated with 10 μM DTG, EVG, TDF, FTC, or three combinations (DTG/TDF/FTC, EVG/TDF/FTC, or TDF/FTC; all ARVs at 10 μM). DTG/TDF/FTC has been the primary cART regimen in use in Botswana since 2016 for all people with HIV, including pregnant women^59^. TDF/FTC and EVG/TDF/FTC are components of Truvada and Stribild, respectively (made by Gilead Sciences, Stribild also features cobicistat).

Treatment with single or combination ARVs for one day did not affect Neuronal Viability, Total Neurite Length, or Neurite Length per Neuron (Fig. 2A-C), although blebbistatin increased Total Neurite Length by 15% (Fig. 2B). By contrast, seven-day ARV and cART exposure caused significant neurotoxicity. Neuronal Viability was significantly reduced by 20% by 10 μM DTG, 80% by 10 μM EVG, 30% by 10μM TDF, 70% by DTG/TDF/FTC, 85% by EVG/TDF/FTC, and 30% by TDF/FTC (Fig. 2D). We used the Tukey’s multiple comparisons test to test for significant differences between data from all test conditions and the DMSO control and between single ARVs to the ARV combinations (see Supplemental Data for full results). Comparing ARV combinations to their components, TDF/FTC treatment led to similar Neuronal Viability to that of TDF alone, suggesting FTC does not affect TDF toxicity (Table S1). EVG/TDF/FTC treatment also led to similar Neuronal Viability to that of EVG alone, suggesting that TDF and FTC do not affect EVG toxicity. By contrast, DTG/TDF/FTC treatment significantly reduced Neuronal Viability compared to either DTG alone or TDF/FTC (p<0.001 for both comparisons), suggesting additive toxicity by these ARVs.

**Figure 2.**
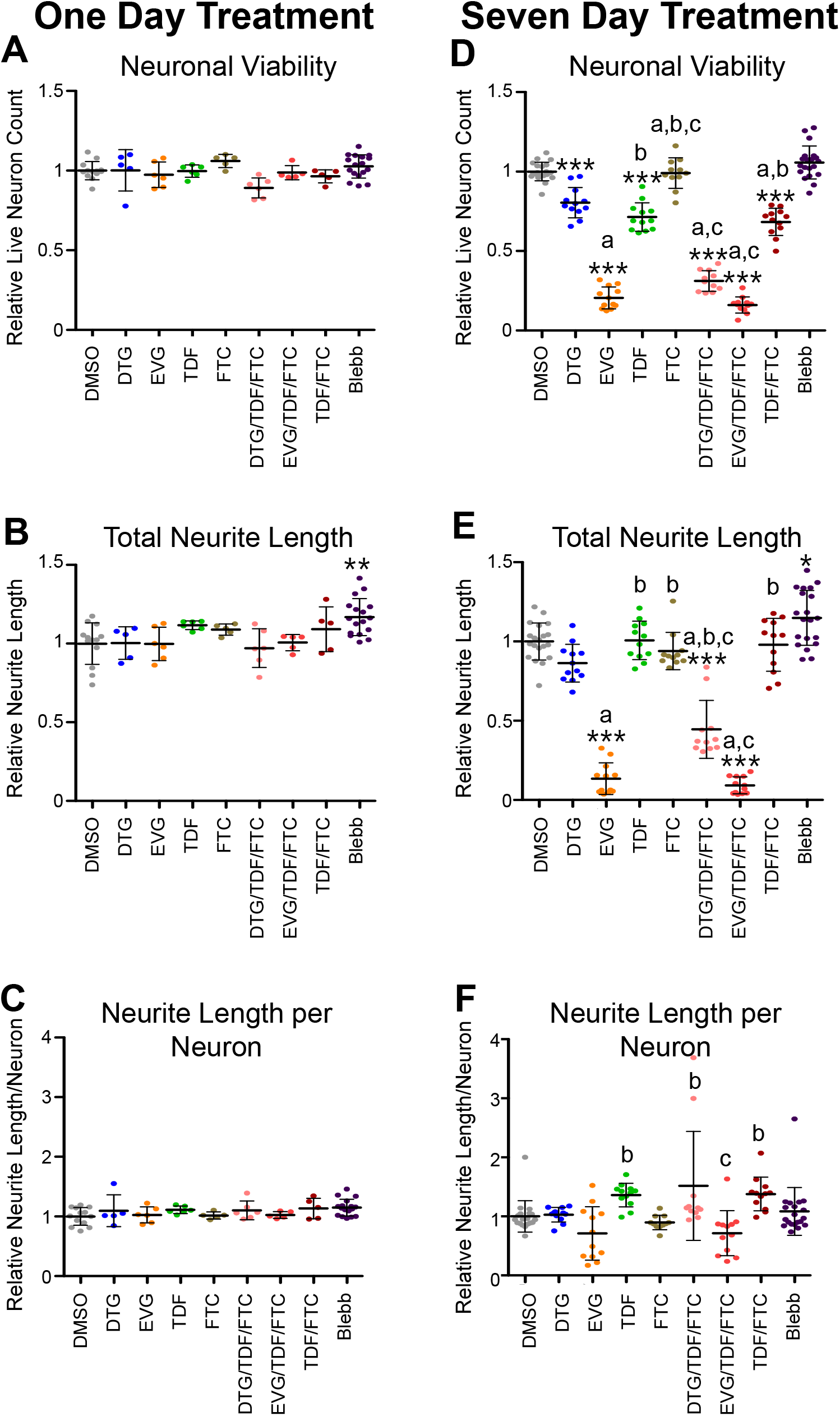
Single and combined ARVs decrease viability and neurite length in hiPSC-neurons after seven days, but not one day of exposure. hiPSC-neurons were treated for one (left) or seven days (right) with DMSO alone, 25 μM blebbistatin, single ARVs, or combinations of ARVs (each at 10 μM). (A, D) Neuronal viability as defined by the number of live neuron nuclei in each well relative to the DMSO control mean. (B, E) Total neurite length in μm per cm^2^ image area in each well relative to the DMSO control mean. (C, F) Total neurite length per live neuron nucleus in each well relative to the DMSO control mean. Each dot represents the average of each measurement from nine images per well. One-day treatment DMSO and blebbistatin: n = 18 wells; ARVs: n = 6 wells. Seven-day treatment DMSO and blebbistatin: n = 36 wells; ARVs: n = 12 wells from two experiments. Bars represent mean ± standard deviation. Statistics performed with one-way ANOVA followed by Tukey’s multiple comparisons test. * p < 0.05, ** p < 0.01, *** p < 0.001. Significant differences between DMSO and ARV or blebbistatin treatments are indicated on each graph. Significant differences between DTG and other ARVs/combinations are indicated with “a”, significant differences between EVG and other ARVs/combinations are indicated with “b”, and significant differences between TDF and other ARVs/combinations are indicated with “c”. Results of other Tukey’s comparisons are reported in Figures S1-S3.

Seven-day ARV exposure also significantly reduced Total Neurite Length for hiPSC-neurons treated with 10 μM EVG (85% reduction), DTG/TDF/FTC (55%), and EVG/TDF/FTC (90%) (Fig. 2E). DTG/FTC/TDF treatment led to a 50% lower Total Neurite Length than DTG alone (p<0.001), while EVG/TDF/FTC treatment led to a similar neurite length to that of EVG alone. Neurite lengths were similar to DMSO for hiPSC-neurons treated with TDF, FTC, or TDF/FTC. Neurite Length per Neuron did not change significantly with any ARV treatment, but there was increased variability in wells treated with EVG, DTG/TDF/FTC, and EVG/TDF/FTC (Fig. 2F). See Tables S2 and S3 for the results of comparisons between each condition for Total Neurite Length and Neurite Length per Neuron.

For both Neuronal Viability and Total Neurite Length, EVG displayed dominant neurotoxic effects in combination treatments (EVG alone reduced Neuronal Viability and Total Neurite Length to the same degree as EVG/TDF/FTC), while DTG and TDF had additive effects (DTG/TDF/FTC reduced Neuronal Viability and Total Neurite Length to a greater extent than either DTG or TDF/FTC).

### HCA to characterize HIV ARV effects on synapses in hiPSC-neurons

To measure the effects of ARV treatments on hiPSC-neuron synapses, we used a CyteSeer image analysis algorithm that identifies synapses as SV2 (presynaptic) and PSD95 (postsynaptic) puncta near each other and Tuj-1-positive neurites. We defined Synapse Density as the number of synapses per cm^2^ of the imaging area and Synapses/Neurite Length as the number of synapses per Total Neurite Length (in microns). Figure 3A shows representative images of SV2 (left) and PSD95 (center) staining of hiPSC-neurons exposed to ARVs for seven days and the live neuronal nuclei (green), neurites (cyan), and synapses (magenta) identified by CyteSeer (right). hiPSC-neurons treated with 10 μM EVG had fewer synapses, with smaller effects for the other ARVs. Effects on hiPSC-neuron synapses appeared at lower doses and for more ARVs than the effects on neuronal viability and neurite length. Synapse Density was significantly reduced by 1 μM DTG (30%), 0.1 μM (20%) and 10 μM (75%) EVG, 1 μM (30%) and 10 μM (40%) TDF, 1 μM (20%) and 10 μM (30%) FTC, and blebbistatin (30%) (Fig. 3B). DTG significantly reduced the Synapses/Neurite Length by 20% at 0.1 μM and 30% at 1 μM but had no effect at 10 μM (Fig. 3C). EVG decreased the Synapses/Neurite Length by 25% at 0.1 μM, had no effect at 1 μM, and increased the Synapses/Neurite Length by 20% at 10 μM. TDF and FTC reduced the Synapses/Neurite Length at all tested concentrations (20%, 25%, and 20% for 0.1, 1, and 10 μM TDF and 20%, 20%, and 30% for 0.1, 1, and 10 μM FTC). Blebbistatin reduced the Synapses/Neurite Length by 20%.

**Figure 3.**
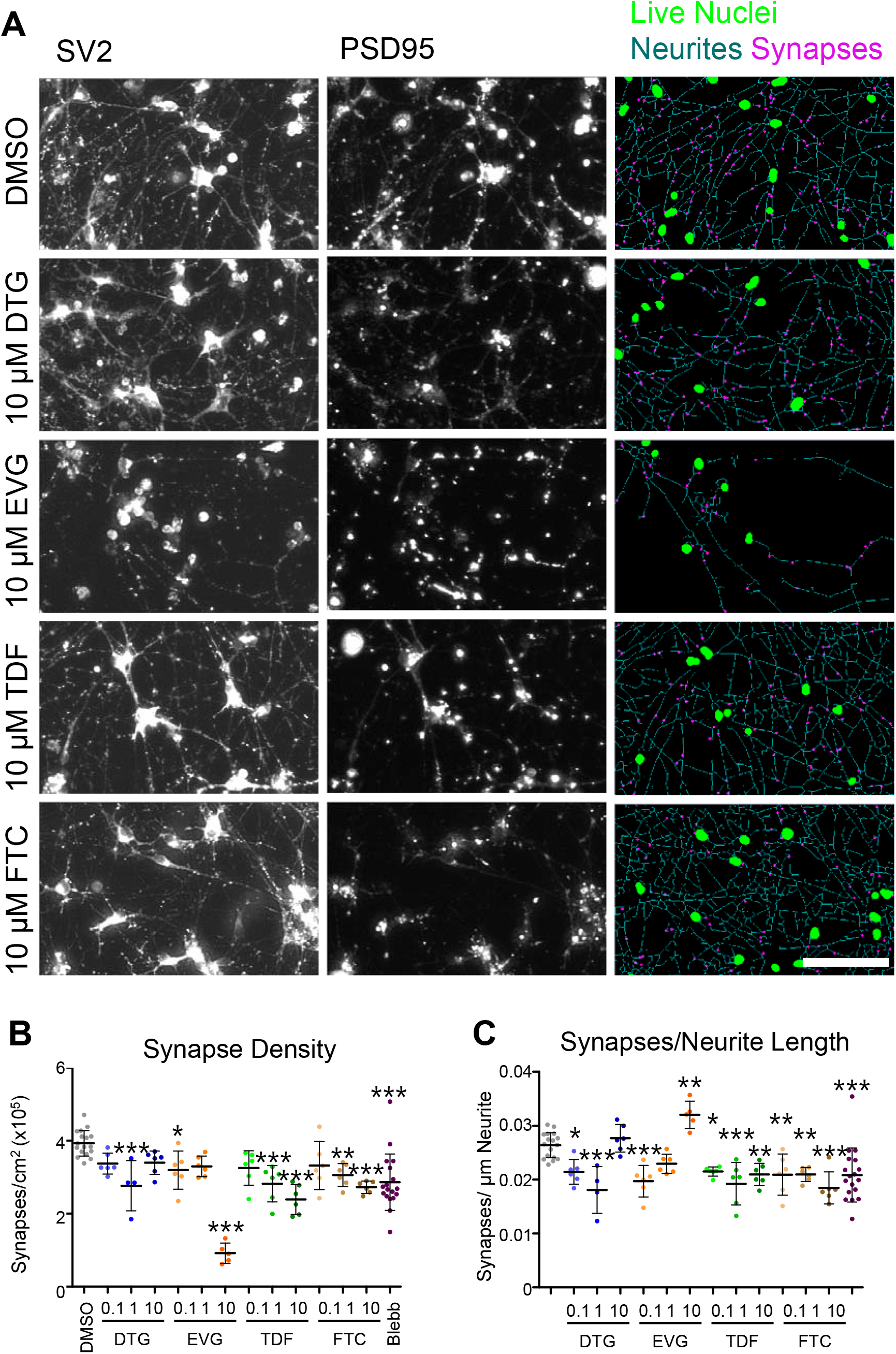
Single ARVs affect synapse density in hiPSC-neurons in a dose-dependent manner. (A) Representative images of hiPSC-neurons from the same experiment in Figure 1 (seven-day ARV treatment) stained for SV2 (presynaptic marker, left) and PSD95 (postsynaptic marker, center). Right images show live neuron nuclei (green), neurites (cyan), and synapses (magenta) identified by CyteSeer. Scale bar = 50 μm. (B) Synapse Density (synapses per cm^2^ imaging area) for each condition. (C) Synapses/Neurite Length (synapses per μm neurite length) for each condition. Each dot represents the average of each measurement from nine images per well. DMSO and blebbistatin: n = 18 wells; ARVs: n = 6 wells. Bars represent mean ± standard deviation. Statistics performed with one-way ANOVA followed by Dunnett’s multiple comparisons test. * p < 0.05, ** p < 0.01, *** p < 0.001.

In experiments comparing 10 μM doses of single ARVs to ARV combinations, one-day treatments did not affect the Synapse Density or Synapses/Neurite Length, although blebbistatin increased the Synapse Density by 20% (Fig 4A, B). By contrast, the Synapse Density was significantly reduced by seven-day treatment with EVG (90%), DTG/TDF/FTC (60%), EVG/TDF/FTC (90%), and TDF/FTC (20%) (Fig. 4C). DTG/FTC/TDF treatment led to a 50% lower Synapse Density than that of DTG alone (p<0.001, Tukey’s), while EVG/TDF/FTC treatment led to a similar Synapse Density to that of EVG alone. TDF/FTC treatment reduced the Synapses/Neurite Length by 20% (vs. DMSO, p<0.001), but this parameter was not affected by other ARV treatments (Fig. 4D). See Tables S4 and S5 for the results of comparisons between each condition for Synapse Density and Synapses/Neurite Length.

**Figure 4.**
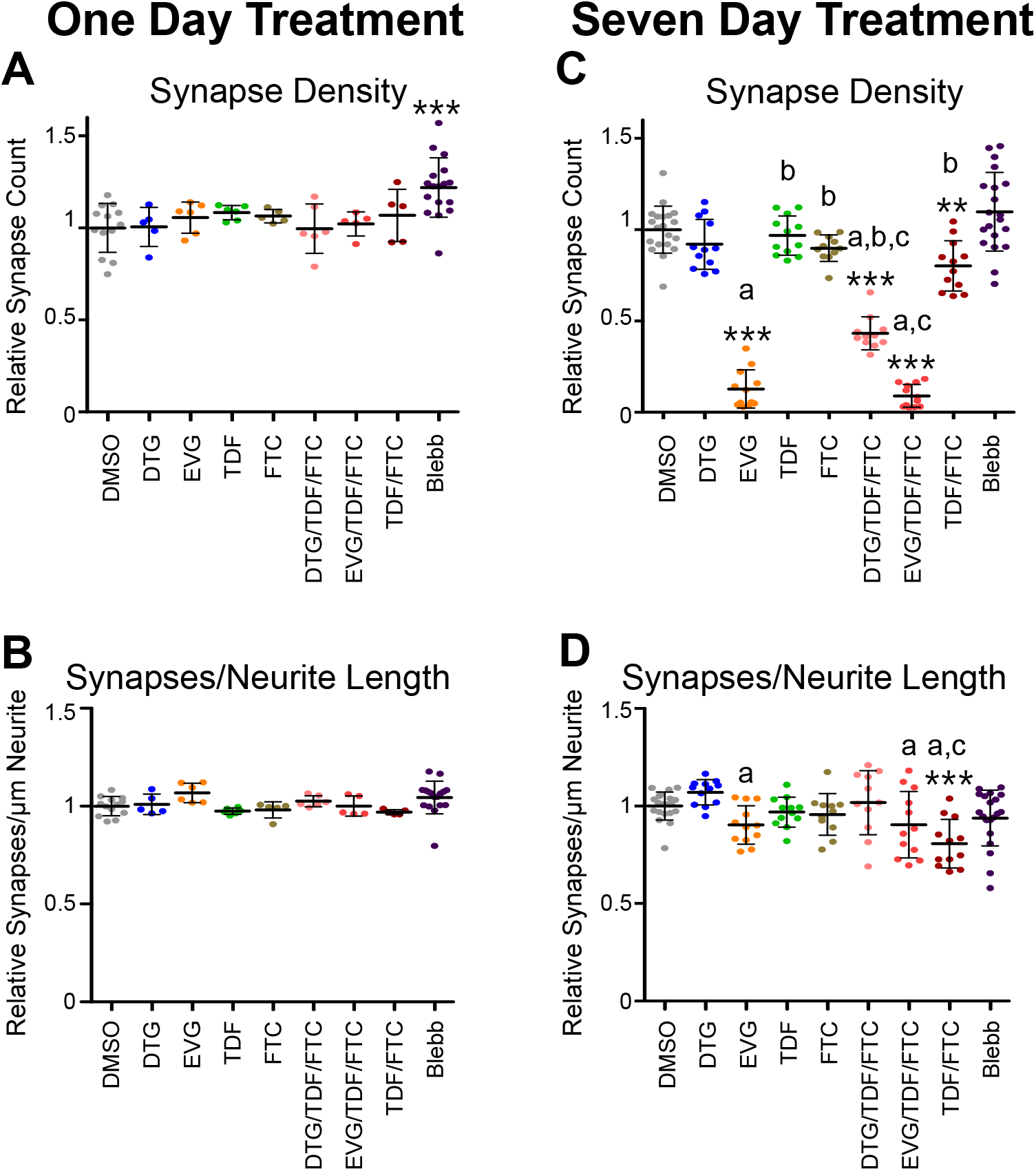
Single and combined ARVs affect synapse density of hiPSC-neurons after seven days, but not one day of exposure. hiPSC-neurons were treated for one or seven days with DMSO alone, 25 μM blebbistatin, single ARVs, or combinations of ARVs (each at 10 μM). (A, C) Synapse Density (synapses per cm^2^ image area) in each well relative to the DMSO control mean. (B, D) Synapses/Neurite Length (synapses per μm neurite length) in each well relative to the DMSO control mean. Each dot represents the average of each measurement from nine images per well. One day treatment DMSO and blebbistatin: n = 18 wells; ARVs: n = 6 wells. Seven day treatment DMSO and blebbistatin: n = 36 wells; ARVs: n = 12 wells from two experiments. Bars represent mean ± standard deviation. Statistics performed with one-way ANOVA followed by Tukey’s multiple comparisons test. * p < 0.05, ** p < 0.01, *** p < 0.001. Significant differences between DMSO and ARV or blebbistatin treatments are indicated on each graph. Significant differences between DTG and other ARVs/combinations are indicated with “a”, significant differences between EVG and other ARVs/combinations are indicated with “b”, and significant differences between TDF and other ARVs/combinations are indicated with “c”. Results of other Tukey’s comparisons are reported in Figures S4 and S5.

### KIC to characterize HIV ARV effects on intracellular calcium transients in hiPSC-neurons

Neurons exhibit action potential-dependent and -independent peaks in intracellular calcium concentration that induce molecular and structural changes within neurons through calcium-sensitive effectors^60–62^. Dysregulation of neuronal intracellular calcium concentration and signaling occurs in aging, traumatic brain injury, and neurodegenerative diseases^63^. To test ARVs for effects on neuronal calcium transients, we treated hiPSC-neurons with 10 μM of each ARV alone or in combination for one or seven days. We then loaded the hiPSC-neurons with Hoechst and the calcium indicator Rhod-4 and assayed the cells for calcium activity. For each well, we collected a single image in the nuclear channel and a digital movie in the Rhod-4 channel at four frames per second for two minutes. We then used a CyteSeer KIC analysis algorithm originally developed for quantifying calcium transients in cardiac myocytes^64^ and adapted to quantify neuronal calcium transients^50^. The algorithm measured the Rhod-4 signal in the soma associated with each live neuron nucleus at each frame, enabling detection of each calcium transient that occurred in each neuron during the recording period. The algorithm then calculated parameters including the percent of neurons with calcium transients and frequency and amplitude of the transients.

hiPSC-neurons exposed to DMSO alone for 7 days displayed between 0 and 50 calcium transients that were typically 5 to 30 seconds in duration during the two-minute recording periods (Fig. 5A). hiPSC-neurons exposed to TDF, FTC, and TDF/FTC for 7 days displayed similar calcium transient activity to those treated with DMSO alone (Fig. 5D, E, H), while neurons treated with DTG or DTG/TDF/FTC had less calcium transient activity compared to DMSO (Fig. 5B, F) and neurons treated with EVG or EVG/TDF/FTC were virtually inactive (Fig. 5C, G).

**Figure 5.**
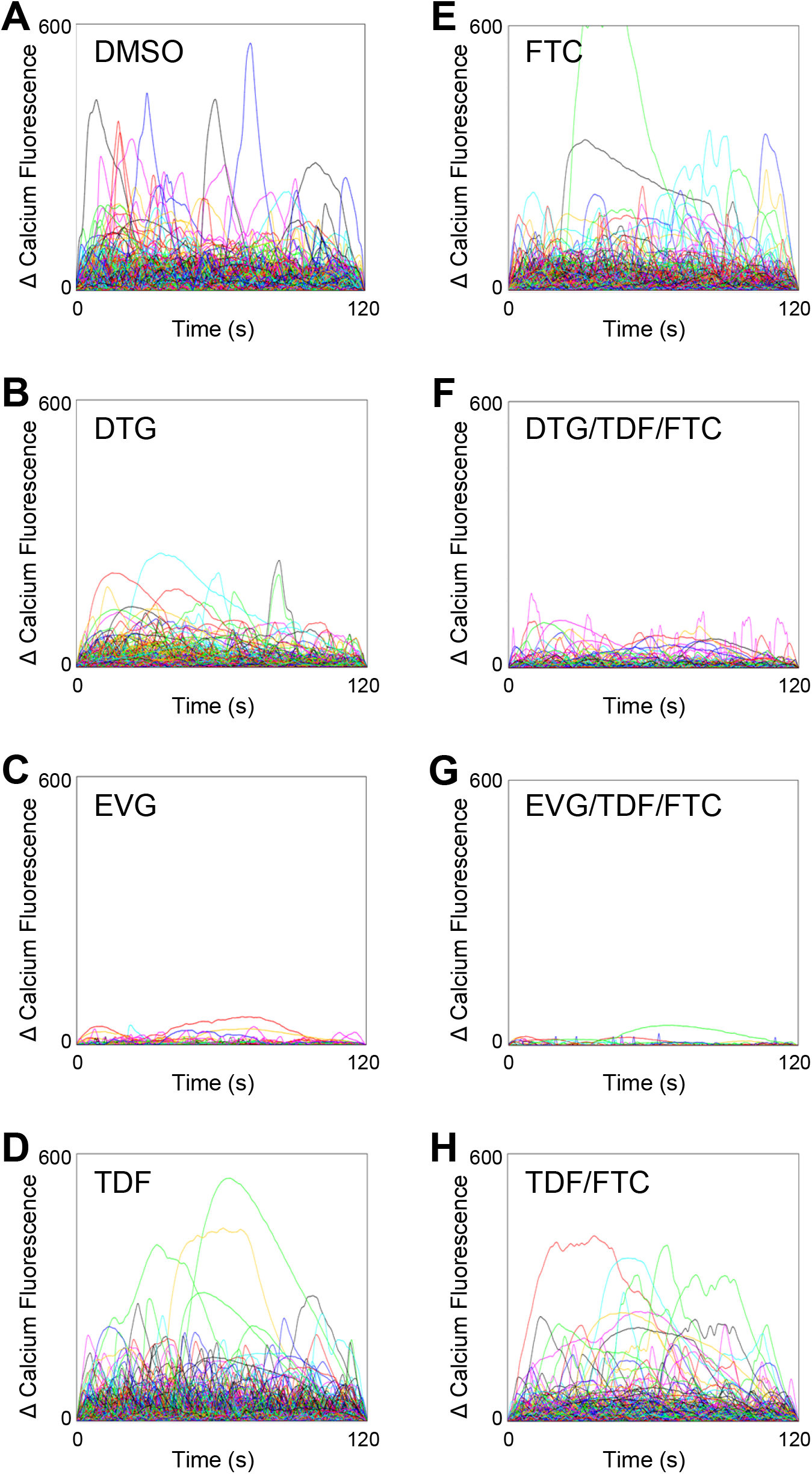
Effect of ARVs on calcium transients in hiPSC-neurons. hiPSC-neurons were treated for seven days with DMSO alone, single ARVs, or combinations of ARVs (each at 10 μM). (A-H) Traces showing transient increases in calcium fluorescence relative to baseline. Each graph contains one trace for each active neuron in each condition with detectable increases in calcium fluorescence. DMSO: 1034 cells; DTG: 716 cells; EVG: 31 cells; TDF: 729 cells; FTC: 1003 cells; DTG/TDF/FTC: 134 cells; EVG/TDF/FTC: 17 cells; TDF/FTC: 539 cells. Six wells per condition.

Quantification of the calcium transient activity showed no significant effects after one day of ARV exposure (Fig. 6A-C). After seven days of ARV exposure, the percent active hiPSC-neurons was reduced by 30% by DTG, 90% by EVG, 70% by DTG/TDF/FTC, 95% by EVG/TDF/FTC, and 25% by TDF/FTC (Fig. 6D). DTG/TDF/FTC treatment led to 60% lower percent activity than DTG alone (p<0.001), while EVG/TDF/FTC treatment led to a similar effect on percent activity to that of EVG alone. The event frequency (Fig. 6E) was significantly reduced by EVG (40%) and by EVG/TDF/FTC (55%), but not by any of the other treatments (while event frequency for DTG/TDF/FTC was lower than for DMSO, this difference did not achieve statistical significance). The mean peak amplitude was also significantly reduced by 10 μM EVG (55%) and EVG/TDF/FTC (50%), but not by any of the other treatments. See Tables S6-8 for the results of comparisons between each condition for percent activity, event frequency, and mean peak amplitude.

**Figure 6.**
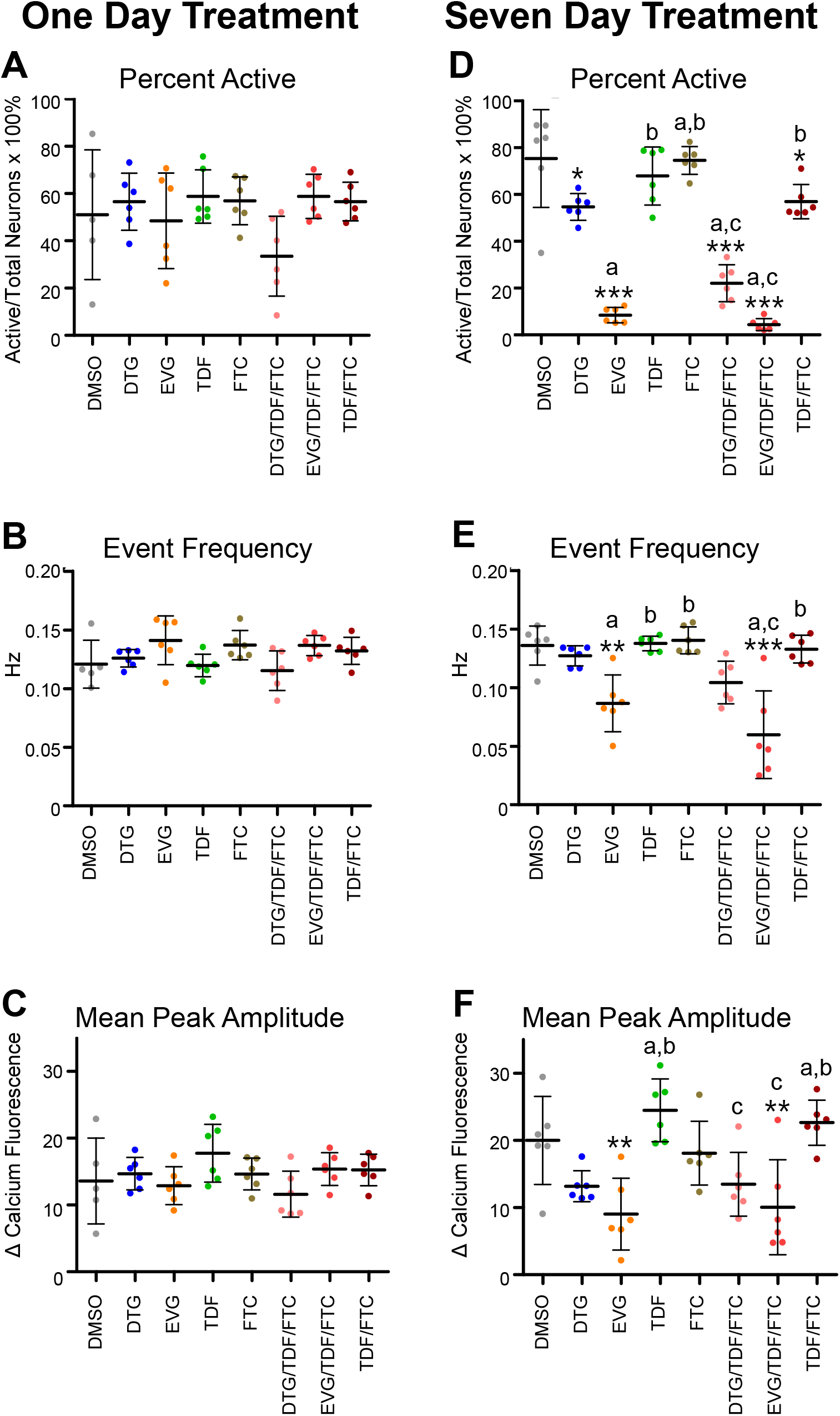
Quantification of the effects of ARVs on calcium activity in hiPSC-neurons. hiPSC-neurons were treated for one or seven days with DMSO alone, single ARVs, or combinations of ARVs (each at 10 μM). (A, D) Percent of live neurons that are active with detectable calcium transients in each condition. (B, E) The mean event frequency of calcium transients of all active neurons in each well. (C, F) The mean of mean calcium peak amplitudes of all active neurons in each well. Six wells per condition. Bars represent mean ± standard deviation. Statistics performed with one-way ANOVA followed by Tukey’s multiple comparisons test. * p < 0.05, ** p < 0.01, *** p < 0.001 vs. DMSO. Significant differences between DTG and other ARVs/combinations are indicated with “a”, significant differences between EVG and other ARVs/combinations are indicated with “b”, and significant differences between TDF and other ARVs/combinations are indicated with “c”. Results of other Tukey’s comparisons are reported in Figures S6-S8.

### Microscopic imaging of the epigenetic landscape (MIEL) analysis to characterize HIV ARV effects on hNPC epigenetic signature

To test for ARV effects on neurogenesis, we treated hNPCs with 0.1,1.0, or 10 μM of each ARV for three days. After treatment with 0.1, 1.0, or 10 μM DTG, EVG, or FTC, hNPC viability was similar to DMSO controls. Treatment with 10 μM TDF, however, significantly reduced hNPC viability by 30% (Fig. 7A). hNPC viability was also reduced by 3 μM SAHA (45%), a histone deacetylase inhibitor, and by 10 μM GSK343 (50%), a histone methyltransferase inhibitor. hNPC viability was not affected by 0.3 μM JQ1, which inhibits the interaction of bromodomain-containing proteins with acetylated histones, or by 1 μM tofacitinib, which inhibits JAK activity.

**Figure 7.**
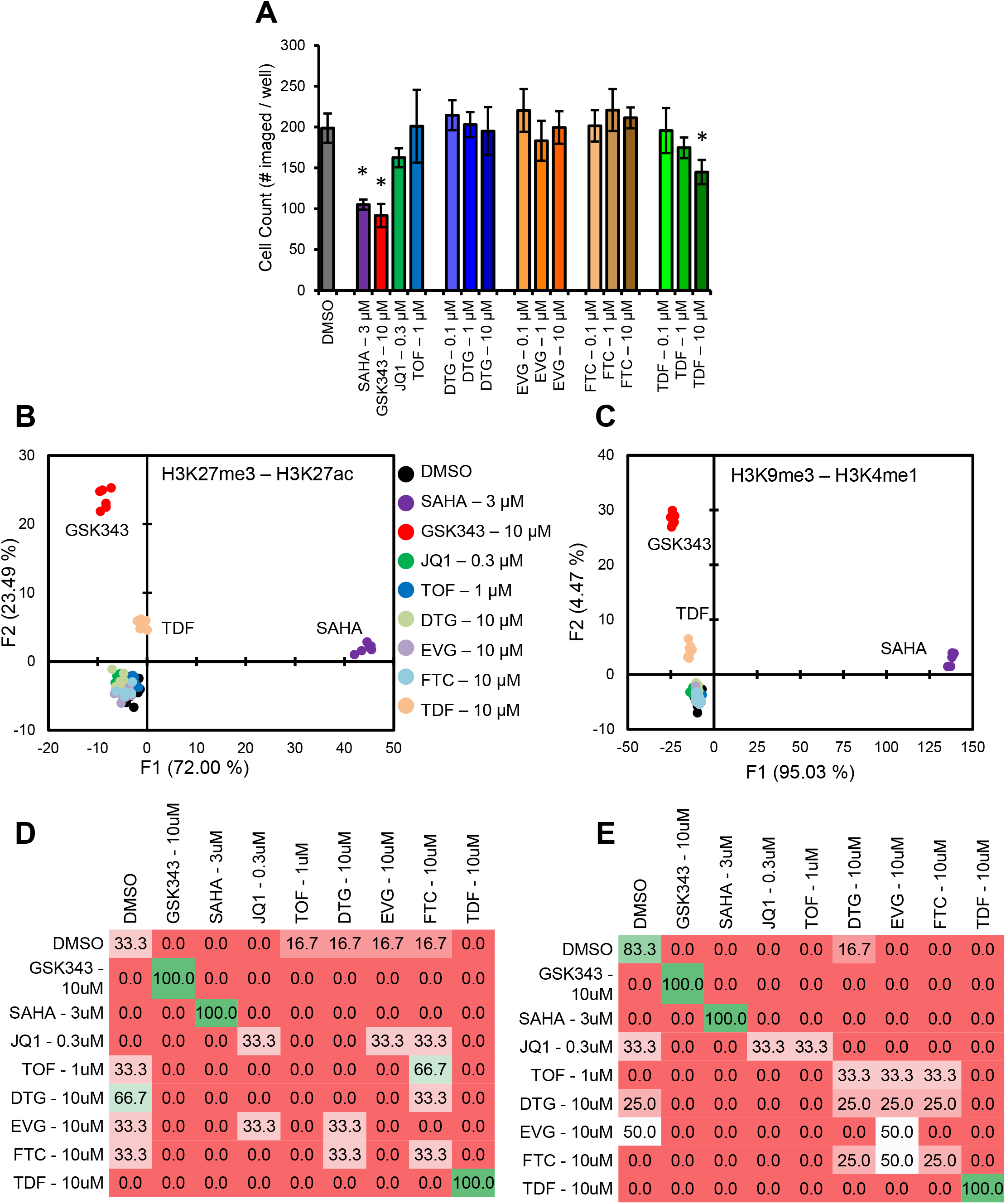
TDF affects the viability and epigenetic signature of hNPCs. Fetal hNPCs were treated with DMSO alone, compounds with known epigenetic effects (SAHA, GSK343, JQ1, or TOF), or ARVs at the indicated concentrations. (A) Count of hNPCs per well for each condition following treatment. Bars represent mean ± standard deviation. Statistics performed with one-way ANOVA followed by Dunnett’s multiple comparisons test. * p < 0.05. (B-E) Quadratic discriminant analysis using texture features derived from images of hNPCs treated with the indicated compounds and immunostained for H3K27me3 and H3K27ac (B, D) or H3K9me3 and H3K4me1 (C, E). (B, C) Scatter plots showing the first two texture-derived discriminant factors for each condition. (D, E) Confusion matrices showing the results of the discriminant analysis. Numbers represent the percent of wells classified correctly (on the diagonal) and incorrectly (off the diagonal). N = 6 wells per condition.

To analyze ARV effects on hNPC epigenetics, we immunolabeled hNPCs for two sets of histone modifications: H3K27me3 (condensed chromatin) and H3K27ac (active enhancers) or H3K9me3 (condensed chromatin) and H3K4me1 (primed enhancers). We then scanned the cells using Vala’s IC200 Image Cytometer and analyzed the images to identify the multivariate epigenetic signatures of each histone modification in each condition. We characterized the epigenetic signatures using texture features^56,65,66^ rather than intensities and morphologies to reduce culturing and immunolabeling artifacts. Quadratic discriminant analysis reports changes in the epigenetic signatures in response to each treatment. SAHA and GSK43, which decreased hNPC viability, also significantly affected the H3K27me3/H3K27ac (Fig. 7B) and the H3K9me3/H3K4me1 (Fig. 7C) epigenetic signatures, with discriminant analysis separating samples treated with these compounds from DMSO- and ARV-treated samples. Of the ARVs tested, only 10 μM TDF altered the epigenetic signature, with TDF-treated samples clustering together and separately from DMSO-, SAHA-, and GSK343-treated samples in both H3K27me3/H3K27ac and H3K9me3/HeK4me1 plots (Fig. 7B, C).

Confusion matrices report the ability of the multiparametric discriminant analysis to correctly classify each sample with other samples that received the same treatment compared to other treatments (Fig. 7D, E)^56^. The confusion matrix for both H3K27me3/H3K27ac and H3K9me3/H3K4me1 show that classification occurred correctly 100% of the time for samples treated with 10 μM GSK343, 3 μM SAHA, and 10 μM TDF. This classification accuracy supports our conclusion that these three compounds are epigenotoxic and significantly alter the hNPC epigenetic landscape. Samples treated with DMSO, DTG, EVG, FTC, JQ1, or TOF, which clustered together after discriminant analysis, were correctly classified less than 100% of the time.

The results from the MIEL assay demonstrate that TDF, but not DTG, EVG, or FTC, reduced viability and altered the pattern of the H3K27me3, H3K27ac, H3K9me3, and H3K4me1 epigenetic histone modifications in hNPCs.

## Discussion

In this study, we developed HCA and KIC methods to quantify neurotoxic and neurodevelopmental effects of HIV ARVs in hiPSC-neurons and human neural precursor cells (Fig. 8). Loss of function and/or cell death in excitatory neurons resulting from ARV neurotoxicity could contribute to HIV-associated neurocognitive disorders (HAND), which currently affect about half of people with HIV^17,18^. Because people with HIV develop non-AIDs-related health conditions, including cognitive decline, at an earlier age than the general population^67–69^, HAND is likely to increase in incidence and severity as the percentage of people with HIV over 50 years old increases^70^. ARV exposure during fetal development, childhood, and adolescence may also increase HAND incidence by affecting neurogenesis at key stages in CNS development. Human in vitro systems are needed to identify and mitigate ARV neurotoxicity and neurodevelopmental effects. The methods used in this study feature automated digital microscopy and analysis for cell plated in 384-well dishes, a format that facilitates higher throughput testing of ARVs at multiple concentrations and combinations, as well as testing of potential HAND therapeutics that may mitigate ARV neurotoxicity.

**Figure 8.**
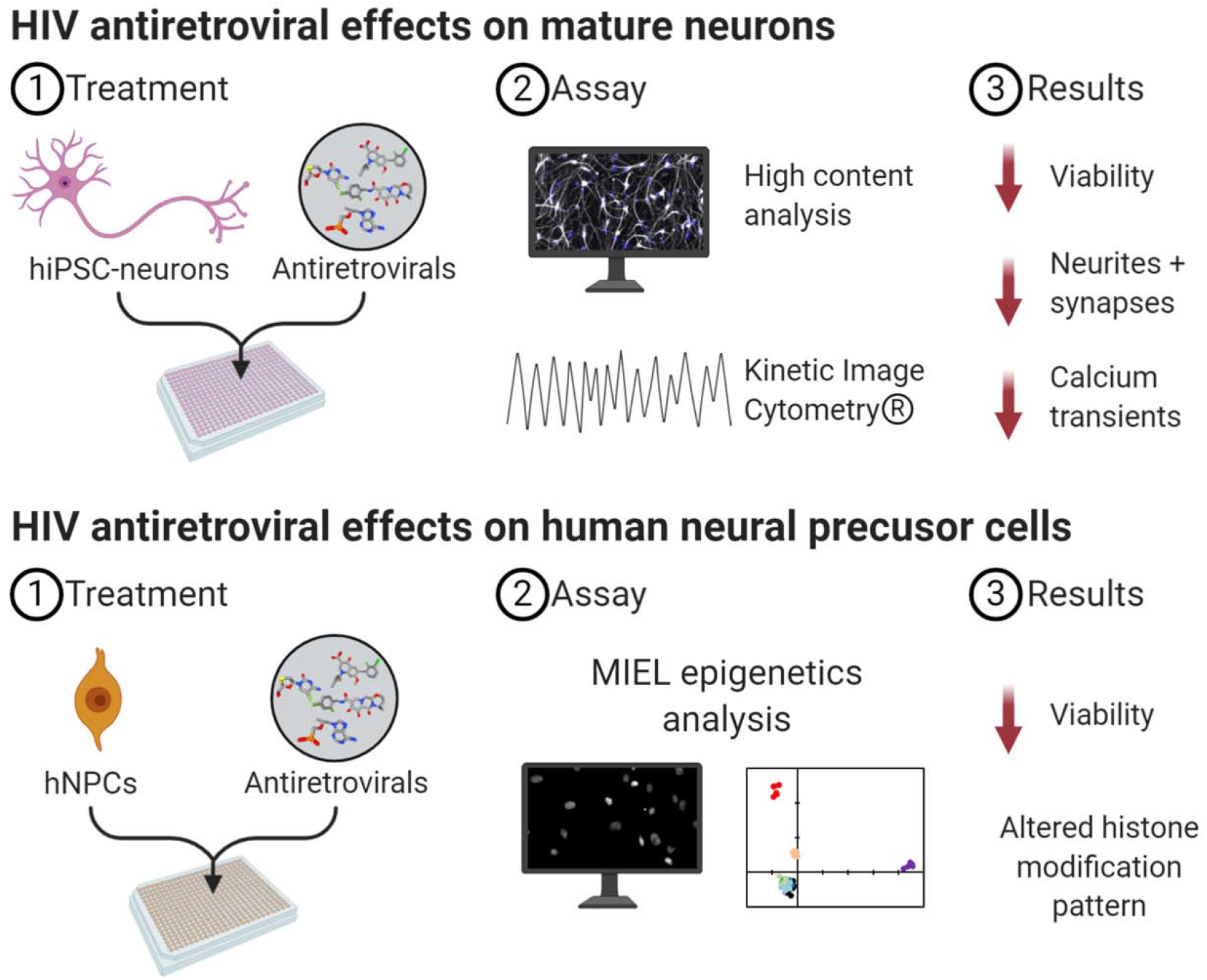
HCA, KIC, and MIEL to identify diverse neurotoxic effects of HIV antiretrovirals. (Top) Outline of our methods to identify HIV antiretroviral effects in hiPSC-neurons. After culture in imaging-quality 384-well plates, hiPSC-neurons are exposed to ARVs alone or in combination for one or seven days. hiPSC-neurons are then fixed and stained for high content analysis or imaged live for KIC of intracellular calcium transients. These assays test if ARVs alter hiPSC-neuron viability, neurite length, synapse density, and/or calcium transient activity. (Bottom) Outline of our methods to identify ARV effects in hNPCs. After culture in imaging-quality 384-well plates, hNPCs are exposed to ARVs for three days. hNPCs are then fixed and stained for microscopic imaging of the epigenetic landscape (MIEL) analysis. This assay tests if ARVs affect hNPC viability and/or change the histone modification pattern within the nuclei. Figure made with Biorender.com.

Many previous in vitro studies of ARV neurotoxicity have been conducted with primary embryonic rat neurons. Exposure of these cells to 10 μM of the integrase inhibitor EVG for four days, but not two days, significantly reduced the number of MAP2+ neurons^28^, while 0.1 and 1 μM EVG were not neurotoxic. In this study, we found that seven-day exposure (but not one-day exposure) to 10 μM EVG reduced hiPSC-neuron viability, decreased neurite length and synapse density, and inhibited intracellular calcium transients, confirming the neurotoxicity of this compound in a human neuron model. We also observed milder toxic effects after treatment with 10 μM of the integrase inhibitor DTG and 10 μM of the nucleoside/nucleotide reverse transcriptase inhibitor TDF. The nucleoside/nucleotide reverse transcriptase inhibitor FTC, which forms a base for front-line recommended cART regimens in the US, was relatively non-toxic in our hiPSC-neuron model.

Our data suggest that plasma, CSF, and brain concentrations of DTG, EVG, and TDF need to be managed to control HIV replication while minimizing ARV neurotoxicity. Efforts to increase the CNS penetration effectiveness (CPE) of ARVs such as EVG^71^ and protease inhibitors^72^ may decrease the CNS viral load but may also increase ARV neurotoxicity. Neuroimaging studies have shown that HAND patients on cART have locally compromised blood brain barriers^73^, which could increase ARV levels in the brain, even for ARVs with low CPE scores. In postmortem brains from HIV-positive people who died while on cART, higher ARV levels were associated with worse antemortem cognitive performance^74^. This association could result from HIV-induced systemic inflammation that leads to blood brain barrier compromise or from increased ARV neurotoxicity. Our human in vitro assays enable direct testing of ARVs for neurotoxicity, which is likely to be relevant to HAND pathology.

While our assay used hiPSC-neurons in isolation, neurons coexist with glial cells like microglia and astrocytes in vivo, which impact their survival and function^75^ and play key roles in neuroinflammation and neurodegenerative diseases such as Alzheimer’s^76,77^. A recently developed in vitro model enables testing for ARV neurotoxicity on co-cultured hiPSC-neurons, - microglia, and -astrocytes^78^ in the presence or absence of HIV infection. In this system, efavirenz, a nucleoside/nucleotide reverse transcriptase inhibitor linked to neuropsychiatric and cognitive effects^79–81^, increased inflammation and reduced phagocytosis in the hiPSC-microglia. Our hiPSC-neuron assay could be similarly expanded to include hiPSC-microglia and -astrocytes to test whether glia contribute to or protect against ARV-induced neuronal toxicity. This expanded system will also enable testing of the efficacy of ARVs to inhibit HIV infection of human microglia and the resulting inflammation^82,83^.

Our assay of ARV effects in hNPCs indicated that TDF reduces viability in this cell type. TDF and zidovidine (AZT) reduce the viability of murine NPCs^84,85^, but to our knowledge this is the first report of ARV effects on human NPCs. TDF also altered the distribution of histone modifications in hNPCs, which may indicate that TDF affects hNPC differentiation. Because ARVs are prescribed as combinations of several drugs, it will be important to investigate such combinations for their epigenotoxic activity. DNA methylation, another epigenetic modification, has recently been reported to be altered in HIV+ children receiving cART^86^. Given the importance of neurogenesis in neurodevelopment and maintenance of cognitive function during aging, it is critical to evaluate existing and candidate ARVs for effects on hNPC viability and epigenetics.

In summary, we have developed and applied methods for the high throughput testing of HIV ARVs on hiPSC-neurons that model excitatory neurons of the human brain. Similar HCA and KIC methods have been applied to quantify neurotoxic effects of the breast cancer therapeutic tamoxifen in primary rat hippocampal neurons^50^, suggesting a broad applicability for these methods for neurotoxicity screening. We have also developed methods to screen for ARV effects on neurogenesis by quantifying the viability and epigenetics of hNPCs. Developing cART regimens that balance the neuroprotective effects of suppressing CNS HIV replication with potential ARV neurotoxicity remains a major challenge in HIV/AIDS treatment and drug discovery^87–89^. The results of the present study represent progress in this direction. Future studies will incorporate human glial cells (hiPSC-microglia and hiPSC-astrocytes) into our assay system to increase its power to detect neurotoxic and neurodevelopmental effects of existing and candidate anti-HIV therapeutics and to identify potential HAND therapeutics that can mitigate these effects.

## Acknowledgements

This study was funded, in part, by grants from the NIH, which include R44ES026268 “Assay of chemicals for Parkinson's toxicity in human iPSC-derived neurons” and R41MH119621 “The Microscopic Imaging of Epigenetic Landscape- NeuroDevelopment (MIEL-ND) assay” and 1R43AG062012-01 “The Alzheimer's Therapeutics Screening Assay: a high-throughput drug-discovery platform utilizing neurons and microglia derived from human induced pluripotent stem cells and Kinetic Image Cytometry”.

---

**Table S1.** Tukey’s multiple comparison test results for Neuronal Viability. Tables show the results of each pairwise comparison for hiPSC-neurons treated for one (left) or seven days (right) with DMSO alone, 25 μM blebbistatin, single ARVs, or combinations of ARVs (each at 10 μM). See Figure 2A, D for graph of results. ns: not significant (p> 0.05) * p < 0.05, ** p < 0.01, *** p < 0.001.

One Day Treatment
| Tukey's Multiple Comparison Test | Mean Difference | Significance |
| --- | --- | --- |
| DMSO vs DTG | -0.002360 | ns |
| DMSO vs EVG | 0.02578 | ns |
| DMSO vs TDF | 0.002801 | ns |
| DMSO vs FTC | -0.06047 | ns |
| DMSO vs DTG TDF FTC | 0.1080 | ns |
| DMSO vs EVG TDF FTC | 0.01235 | ns |
| DMSO vs TDF FTC | 0.03525 | ns |
| DMSO vs Blebb | -0.02633 | ns |
| DTG vs EVG | 0.02814 | ns |
| DTG vs TDF | 0.005161 | ns |
| DTG vs FTC | -0.05811 | ns |
| DTG vs DTG TDF FTC | 0.1104 | ns |
| DTG vs EVG TDF FTC | 0.01471 | ns |
| DTG vs TDF FTC | 0.03761 | ns |
| DTG vs Blebb | -0.02397 | ns |
| EVG vs TDF | -0.02298 | ns |
| EVG vs FTC | -0.08625 | ns |
| EVG vs DTG TDF FTC | 0.08225 | ns |
| EVG vs EVG TDF FTC | -0.01343 | ns |
| EVG vs TDF FTC | 0.009474 | ns |
| EVG vs Blebb | -0.05211 | ns |
| TDF vs FTC | -0.06328 | ns |
| TDF vs DTG TDF FTC | 0.1052 | ns |
| TDF vs EVG TDF FTC | 0.009544 | ns |
| TDF vs TDF FTC | 0.03245 | ns |
| TDF vs Blebb | -0.02913 | ns |
| FTC vs DTG TDF FTC | 0.1685 | ** |
| FTC vs EVG TDF FTC | 0.07282 | ns |
| FTC vs TDF FTC | 0.09573 | ns |
| FTC vs Blebb | 0.03414 | ns |
| DTG TDF FTC vs EVG TDF FTC | -0.09568 | ns |
| DTG TDF FTC vs TDF FTC | -0.07277 | ns |
| DTG TDF FTC vs Blebb | -0.1344 | ** |
| EVG TDF FTC vs TDF FTC | 0.02291 | ns |
| EVG TDF FTC vs Blebb | -0.03868 | ns |
| TDF FTC vs Blebb | -0.06158 | ns |

Seven Day Treatment
| Tukey's Multiple Comparison Test | Mean Difference | Significance |
| --- | --- | --- |
| DMSO vs DTG | 0.1952 | *** |
| DMSO vs EVG | 0.7947 | *** |
| DMSO vs TDF | 0.2858 | *** |
| DMSO vs FTC | 0.009897 | ns |
| DMSO vs DTG TDF FTC | 0.6876 | *** |
| DMSO vs EVG TDF FTC | 0.8388 | *** |
| DMSO vs TDF FTC | 0.3164 | *** |
| DMSO vs Blebb | -0.05650 | ns |
| DTG vs EVG | 0.5994 | *** |
| DTG vs TDF | 0.09056 | ns |
| DTG vs FTC | -0.1853 | *** |
| DTG vs DTG TDF FTC | 0.4923 | *** |
| DTG vs EVG TDF FTC | 0.6436 | *** |
| DTG vs TDF FTC | 0.1212 | * |
| DTG vs Blebb | -0.2517 | *** |
| EVG vs TDF | -0.5089 | *** |
| EVG vs FTC | -0.7848 | *** |
| EVG vs DTG TDF FTC | -0.1071 | ns |
| EVG vs EVG TDF FTC | 0.04412 | ns |
| EVG vs TDF FTC | -0.4783 | *** |
| EVG vs Blebb | -0.8512 | *** |
| TDF vs FTC | -0.2759 | *** |
| TDF vs DTG TDF FTC | 0.4018 | *** |
| TDF vs EVG TDF FTC | 0.5530 | *** |
| TDF vs TDF FTC | 0.03061 | ns |
| TDF vs Blebb | -0.3423 | *** |
| FTC vs DTG TDF FTC | 0.6777 | *** |
| FTC vs EVG TDF FTC | 0.8289 | *** |
| FTC vs TDF FTC | 0.3065 | *** |
| FTC vs Blebb | -0.06640 | ns |
| DTG TDF FTC vs EVG TDF FTC | 0.1512 | *** |
| DTG TDF FTC vs TDF FTC | -0.3712 | *** |
| DTG TDF FTC vs Blebb | -0.7441 | *** |
| EVG TDF FTC vs TDF FTC | -0.5224 | *** |
| EVG TDF FTC vs Blebb | -0.8953 | *** |
| TDF FTC vs Blebb | -0.3729 | *** |

**Table S2.** Tukey’s multiple comparison test results for Total Neurite Length. Tables show the results of each pairwise comparison for hiPSC-neurons treated for one (left) or seven days (right) with DMSO alone, 25 μM blebbistatin, single ARVs, or combinations of ARVs (each at 10 μM). See Figure 2B, E for graph of results. ns: not significant (p> 0.05) * p < 0.05, ** p < 0.01, *** p < 0.001.

One Day Treatment
| Tukey's Multiple Comparison Test | Mean Difference | Significance |
| --- | --- | --- |
| DMSO vs DTG | -0.003525 | ns |
| DMSO vs EVG | 0.001910 | ns |
| DMSO vs TDF | -0.1167 | ns |
| DMSO vs FTC | -0.08899 | ns |
| DMSO vs DTG TDF FTC | 0.02920 | ns |
| DMSO vs EVG TDF FTC | -0.006731 | ns |
| DMSO vs TDF FTC | -0.09129 | ns |
| DMSO vs Blebb | -0.1680 | ** |
| DTG vs EVG | 0.005434 | ns |
| DTG vs TDF | -0.1132 | ns |
| DTG vs FTC | -0.08547 | ns |
| DTG vs DTG TDF FTC | 0.03272 | ns |
| DTG vs EVG TDF FTC | -0.003206 | ns |
| DTG vs TDF FTC | -0.08777 | ns |
| DTG vs Blebb | -0.1645 | ns |
| EVG vs TDF | -0.1186 | ns |
| EVG vs FTC | -0.09090 | ns |
| EVG vs DTG TDF FTC | 0.02729 | ns |
| EVG vs EVG TDF FTC | -0.008641 | ns |
| EVG vs TDF FTC | -0.09320 | ns |
| EVG vs Blebb | -0.1699 | * |
| TDF vs FTC | 0.02772 | ns |
| TDF vs DTG TDF FTC | 0.1459 | ns |
| TDF vs EVG TDF FTC | 0.1100 | ns |
| TDF vs TDF FTC | 0.02542 | ns |
| TDF vs Blebb | -0.05132 | ns |
| FTC vs DTG TDF FTC | 0.1182 | ns |
| FTC vs EVG TDF FTC | 0.08226 | ns |
| FTC vs TDF FTC | -0.002303 | ns |
| FTC vs Blebb | -0.07904 | ns |
| DTG TDF FTC vs EVG TDF FTC | -0.03593 | ns |
| DTG TDF FTC vs TDF FTC | -0.1205 | ns |
| DTG TDF FTC vs Blebb | -0.1972 | ** |
| EVG TDF FTC vs TDF FTC | -0.08456 | ns |
| EVG TDF FTC vs Blebb | -0.1613 | ns |
| TDF FTC vs Blebb | -0.07674 | ns |

Seven Day Treatment
| Tukey's Multiple Comparison Test | Mean Difference | Significance |
| --- | --- | --- |
| DMSO vs DTG | 0.1369 | ns |
| DMSO vs EVG | 0.8657 | *** |
| DMSO vs TDF | -0.006719 | ns |
| DMSO vs FTC | 0.06042 | ns |
| DMSO vs DTG TDF FTC | 0.5543 | *** |
| DMSO vs EVG TDF FTC | 0.9078 | *** |
| DMSO vs TDF FTC | 0.02082 | ns |
| DMSO vs Blebb | -0.1483 | * |
| DTG vs EVG | 0.7288 | *** |
| DTG vs TDF | -0.1436 | ns |
| DTG vs FTC | -0.07650 | ns |
| DTG vs DTG TDF FTC | 0.4174 | *** |
| DTG vs EVG TDF FTC | 0.7708 | *** |
| DTG vs TDF FTC | -0.1161 | ns |
| DTG vs Blebb | -0.2853 | *** |
| EVG vs TDF | -0.8724 | *** |
| EVG vs FTC | -0.8053 | *** |
| EVG vs DTG TDF FTC | -0.3114 | *** |
| EVG vs EVG TDF FTC | 0.04204 | ns |
| EVG vs TDF FTC | -0.8449 | *** |
| EVG vs Blebb | -1.014 | *** |
| TDF vs FTC | 0.06714 | ns |
| TDF vs DTG TDF FTC | 0.5610 | *** |
| TDF vs EVG TDF FTC | 0.9145 | *** |
| TDF vs TDF FTC | 0.02753 | ns |
| TDF vs Blebb | -0.1416 | ns |
| FTC vs DTG TDF FTC | 0.4939 | *** |
| FTC vs EVG TDF FTC | 0.8473 | *** |
| FTC vs TDF FTC | -0.03961 | ns |
| FTC vs Blebb | -0.2088 | ** |
| DTG TDF FTC vs EVG TDF FTC | 0.3535 | *** |
| DTG TDF FTC vs TDF FTC | -0.5335 | *** |
| DTG TDF FTC vs Blebb | -0.7026 | *** |
| EVG TDF FTC vs TDF FTC | -0.8870 | *** |
| EVG TDF FTC vs Blebb | -1.056 | *** |
| TDF FTC vs Blebb | -0.1691 | * |

**Table S3.** Tukey’s multiple comparison test results for Neurite Length per Neuron. Tables show the results of each pairwise comparison for hiPSC-neurons treated for one (left) or seven days (right) with DMSO alone, 25 μM blebbistatin, single ARVs, or combinations of ARVs (each at 10 μM). See Figure 2C, F for graph of results. ns: not significant (p> 0.05) * p < 0.05, ** p < 0.01, *** p < 0.001.

One Day Treatment
| Tukey's Multiple Comparison Test | Mean Difference | Significance |
| --- | --- | --- |
| DMSO vs DTG | -0.09484 | ns |
| DMSO vs EVG | -0.02655 | ns |
| DMSO vs TDF | -0.1107 | ns |
| DMSO vs FTC | -0.01540 | ns |
| DMSO vs DTG TDF FTC | -0.1021 | ns |
| DMSO vs EVG TDF FTC | -0.02473 | ns |
| DMSO vs TDF FTC | -0.1341 | ns |
| DMSO vs Blebb | -0.1484 | ns |
| DTG vs EVG | 0.06828 | ns |
| DTG vs TDF | -0.01582 | ns |
| DTG vs FTC | 0.07943 | ns |
| DTG vs DTG TDF FTC | -0.007285 | ns |
| DTG vs EVG TDF FTC | 0.07010 | ns |
| DTG vs TDF FTC | -0.03929 | ns |
| DTG vs Blebb | -0.05354 | ns |
| EVG vs TDF | -0.08410 | ns |
| EVG vs FTC | 0.01115 | ns |
| EVG vs DTG TDF FTC | -0.07557 | ns |
| EVG vs EVG TDF FTC | 0.001821 | ns |
| EVG vs TDF FTC | -0.1076 | ns |
| EVG vs Blebb | -0.1218 | ns |
| TDF vs FTC | 0.09526 | ns |
| TDF vs DTG TDF FTC | 0.008536 | ns |
| TDF vs EVG TDF FTC | 0.08592 | ns |
| TDF vs TDF FTC | -0.02347 | ns |
| TDF vs Blebb | -0.03772 | ns |
| FTC vs DTG TDF FTC | -0.08672 | ns |
| FTC vs EVG TDF FTC | -0.009331 | ns |
| FTC vs TDF FTC | -0.1187 | ns |
| FTC vs Blebb | -0.1330 | ns |
| DTG TDF FTC vs EVG TDF FTC | 0.07739 | ns |
| DTG TDF FTC vs TDF FTC | -0.03200 | ns |
| DTG TDF FTC vs Blebb | -0.04625 | ns |
| EVG TDF FTC vs TDF FTC | -0.1094 | ns |
| EVG TDF FTC vs Blebb | -0.1236 | ns |
| TDF FTC vs Blebb | -0.01425 | ns |

Seven Day Treatment
| Tukey's Multiple Comparison Test | Mean Difference | Significance |
| --- | --- | --- |
| DMSO vs DTG | -0.02907 | ns |
| DMSO vs EVG | 0.2885 | ns |
| DMSO vs TDF | -0.3622 | ns |
| DMSO vs FTC | 0.1027 | ns |
| DMSO vs DTG TDF FTC | -0.5172 | * |
| DMSO vs EVG TDF FTC | 0.2840 | ns |
| DMSO vs TDF FTC | -0.3790 | ns |
| DMSO vs Blebb | -0.08370 | ns |
| DTG vs EVG | 0.3175 | ns |
| DTG vs TDF | -0.3331 | ns |
| DTG vs FTC | 0.1317 | ns |
| DTG vs DTG TDF FTC | -0.4881 | ns |
| DTG vs EVG TDF FTC | 0.3131 | ns |
| DTG vs TDF FTC | -0.3499 | ns |
| DTG vs Blebb | -0.05463 | ns |
| EVG vs TDF | -0.6507 | ** |
| EVG vs FTC | -0.1858 | ns |
| EVG vs DTG TDF FTC | -0.8056 | *** |
| EVG vs EVG TDF FTC | -0.004459 | ns |
| EVG vs TDF FTC | -0.6675 | ** |
| EVG vs Blebb | -0.3722 | ns |
| TDF vs FTC | 0.4649 | ns |
| TDF vs DTG TDF FTC | -0.1550 | ns |
| TDF vs EVG TDF FTC | 0.6462 | ** |
| TDF vs TDF FTC | -0.01681 | ns |
| TDF vs Blebb | 0.2785 | ns |
| FTC vs DTG TDF FTC | -0.6198 | * |
| FTC vs EVG TDF FTC | 0.1813 | ns |
| FTC vs TDF FTC | -0.4817 | ns |
| FTC vs Blebb | -0.1864 | ns |
| DTG TDF FTC vs EVG TDF FTC | 0.8012 | *** |
| DTG TDF FTC vs TDF FTC | 0.1382 | ns |
| DTG TDF FTC vs Blebb | 0.4335 | ns |
| EVG TDF FTC vs TDF FTC | -0.6630 | ** |
| EVG TDF FTC vs Blebb | -0.3677 | ns |
| TDF FTC vs Blebb | 0.2953 | ns |

**Table S4.** Tukey’s multiple comparison test results for Synapse Density. Tables show the results of each pairwise comparison for hiPSC-neurons treated for one (left) or seven days (right) with DMSO alone, 25 μM blebbistatin, single ARVs, or combinations of ARVs (each at 10 μM). See Figure 4A, C for graph of results. ns: not significant (p> 0.05) * p < 0.05, ** p < 0.01, *** p < 0.001.

One Day Treatment
| Tukey's Multiple Comparison Test | Mean Difference | Significance |
| --- | --- | --- |
| DMSO vs DTG | -0.005561 | ns |
| DMSO vs EVG | -0.05608 | ns |
| DMSO vs TDF | -0.08320 | ns |
| DMSO vs FTC | -0.06424 | ns |
| DMSO vs DTG TDF FTC | 0.004011 | ns |
| DMSO vs EVG TDF FTC | -0.02199 | ns |
| DMSO vs TDF FTC | -0.06814 | ns |
| DMSO vs Blebb | -0.2197 | *** |
| DTG vs EVG | -0.05051 | ns |
| DTG vs TDF | -0.07764 | ns |
| DTG vs FTC | -0.05868 | ns |
| DTG vs DTG TDF FTC | 0.009572 | ns |
| DTG vs EVG TDF FTC | -0.01643 | ns |
| DTG vs TDF FTC | -0.06258 | ns |
| DTG vs Blebb | -0.2142 | * |
| EVG vs TDF | -0.02713 | ns |
| EVG vs FTC | -0.008162 | ns |
| EVG vs DTG TDF FTC | 0.06009 | ns |
| EVG vs EVG TDF FTC | 0.03408 | ns |
| EVG vs TDF FTC | -0.01207 | ns |
| EVG vs Blebb | -0.1636 | ns |
| TDF vs FTC | 0.01897 | ns |
| TDF vs DTG TDF FTC | 0.08722 | ns |
| TDF vs EVG TDF FTC | 0.06121 | ns |
| TDF vs TDF FTC | 0.01506 | ns |
| TDF vs Blebb | -0.1365 | ns |
| FTC vs DTG TDF FTC | 0.06825 | ns |
| FTC vs EVG TDF FTC | 0.04225 | ns |
| FTC vs TDF FTC | -0.003904 | ns |
| FTC vs Blebb | -0.1555 | ns |
| DTG TDF FTC vs EVG TDF FTC | -0.02600 | ns |
| DTG TDF FTC vs TDF FTC | -0.07215 | ns |
| DTG TDF FTC vs Blebb | -0.2237 | ** |
| EVG TDF FTC vs TDF FTC | -0.04615 | ns |
| EVG TDF FTC vs Blebb | -0.1977 | ns |
| TDF FTC vs Blebb | -0.1516 | ns |

Seven Day Treatment
| Tukey's Multiple Comparison Test | Mean Difference | Significance |
| --- | --- | --- |
| DMSO vs DTG | 0.07950 | ns |
| DMSO vs EVG | 0.8726 | *** |
| DMSO vs TDF | 0.03189 | ns |
| DMSO vs FTC | 0.1014 | ns |
| DMSO vs DTG TDF FTC | 0.5669 | *** |
| DMSO vs EVG TDF FTC | 0.9095 | *** |
| DMSO vs TDF FTC | 0.1984 | ** |
| DMSO vs Blebb | -0.09814 | ns |
| DTG vs EVG | 0.7931 | *** |
| DTG vs TDF | -0.04760 | ns |
| DTG vs FTC | 0.02194 | ns |
| DTG vs DTG TDF FTC | 0.4874 | *** |
| DTG vs EVG TDF FTC | 0.8300 | *** |
| DTG vs TDF FTC | 0.1189 | ns |
| DTG vs Blebb | -0.1776 | * |
| EVG vs TDF | -0.8407 | *** |
| EVG vs FTC | -0.7711 | *** |
| EVG vs DTG TDF FTC | -0.3056 | *** |
| EVG vs EVG TDF FTC | 0.03694 | ns |
| EVG vs TDF FTC | -0.6742 | *** |
| EVG vs Blebb | -0.9707 | *** |
| TDF vs FTC | 0.06954 | ns |
| TDF vs DTG TDF FTC | 0.5350 | *** |
| TDF vs EVG TDF FTC | 0.8776 | *** |
| TDF vs TDF FTC | 0.1665 | ns |
| TDF vs Blebb | -0.1300 | ns |
| FTC vs DTG TDF FTC | 0.4655 | *** |
| FTC vs EVG TDF FTC | 0.8081 | *** |
| FTC vs TDF FTC | 0.09693 | ns |
| FTC vs Blebb | -0.1996 | ** |
| DTG TDF FTC vs EVG TDF FTC | 0.3426 | *** |
| DTG TDF FTC vs TDF FTC | -0.3685 | *** |
| DTG TDF FTC vs Blebb | -0.6651 | *** |
| EVG TDF FTC vs TDF FTC | -0.7111 | *** |
| EVG TDF FTC vs Blebb | -1.008 | *** |
| TDF FTC vs Blebb | -0.2965 | *** |

**Table S5.** Tukey’s multiple comparison test results for Synapses/Neurite Length. Tables show the results of each pairwise comparison for hiPSC-neurons treated for one (left) or seven days (right) with DMSO alone, 25 μM blebbistatin, single ARVs, or combinations of ARVs (each at 10 μM). See Figure 4B, D for graph of results. ns: not significant (p> 0.05) * p < 0.05, ** p < 0.01, *** p < 0.001.

One Day Treatment
| Tukey's Multiple Comparison Test | Mean Difference | Significance |
| --- | --- | --- |
| DMSO vs DTG | -0.009334 | ns |
| DMSO vs EVG | -0.06784 | ns |
| DMSO vs TDF | 0.02567 | ns |
| DMSO vs FTC | 0.01931 | ns |
| DMSO vs DTG TDF FTC | -0.02523 | ns |
| DMSO vs EVG TDF FTC | -0.0004753 | ns |
| DMSO vs TDF FTC | 0.03048 | ns |
| DMSO vs Blebb | -0.04413 | ns |
| DTG vs EVG | -0.05851 | ns |
| DTG vs TDF | 0.03500 | ns |
| DTG vs FTC | 0.02864 | ns |
| DTG vs DTG TDF FTC | -0.01589 | ns |
| DTG vs EVG TDF FTC | 0.008858 | ns |
| DTG vs TDF FTC | 0.03981 | ns |
| DTG vs Blebb | -0.03480 | ns |
| EVG vs TDF | 0.09351 | ns |
| EVG vs FTC | 0.08715 | ns |
| EVG vs DTG TDF FTC | 0.04262 | ns |
| EVG vs EVG TDF FTC | 0.06737 | ns |
| EVG vs TDF FTC | 0.09832 | ns |
| EVG vs Blebb | 0.02371 | ns |
| TDF vs FTC | -0.006361 | ns |
| TDF vs DTG TDF FTC | -0.05090 | ns |
| TDF vs EVG TDF FTC | -0.02615 | ns |
| TDF vs TDF FTC | 0.004810 | ns |
| TDF vs Blebb | -0.06980 | ns |
| FTC vs DTG TDF FTC | -0.04454 | ns |
| FTC vs EVG TDF FTC | -0.01979 | ns |
| FTC vs TDF FTC | 0.01117 | ns |
| FTC vs Blebb | -0.06344 | ns |
| DTG TDF FTC vs EVG TDF FTC | 0.02475 | ns |
| DTG TDF FTC vs TDF FTC | 0.05571 | ns |
| DTG TDF FTC vs Blebb | -0.01891 | ns |
| EVG TDF FTC vs TDF FTC | 0.03096 | ns |
| EVG TDF FTC vs Blebb | -0.04366 | ns |
| TDF FTC vs Blebb | -0.07461 | ns |

Seven Day Treatment
| Tukey's Multiple Comparison Test | Mean Difference | Significance |
| --- | --- | --- |
| DMSO vs DTG | -0.07061 | ns |
| DMSO vs EVG | 0.09677 | ns |
| DMSO vs TDF | 0.03095 | ns |
| DMSO vs FTC | 0.04279 | ns |
| DMSO vs DTG TDF FTC | -0.01777 | ns |
| DMSO vs EVG TDF FTC | 0.09549 | ns |
| DMSO vs TDF FTC | 0.1925 | *** |
| DMSO vs Blebb | 0.06131 | ns |
| DTG vs EVG | 0.1674 | * |
| DTG vs TDF | 0.1016 | ns |
| DTG vs FTC | 0.1134 | ns |
| DTG vs DTG TDF FTC | 0.05284 | ns |
| DTG vs EVG TDF FTC | 0.1661 | * |
| DTG vs TDF FTC | 0.2632 | *** |
| DTG vs Blebb | 0.1319 | ns |
| EVG vs TDF | -0.06582 | ns |
| EVG vs FTC | -0.05398 | ns |
| EVG vs DTG TDF FTC | -0.1145 | ns |
| EVG vs EVG TDF FTC | -0.001282 | ns |
| EVG vs TDF FTC | 0.09577 | ns |
| EVG vs Blebb | -0.03546 | ns |
| TDF vs FTC | 0.01184 | ns |
| TDF vs DTG TDF FTC | -0.04872 | ns |
| TDF vs EVG TDF FTC | 0.06454 | ns |
| TDF vs TDF FTC | 0.1616 | * |
| TDF vs Blebb | 0.03036 | ns |
| FTC vs DTG TDF FTC | -0.06056 | ns |
| FTC vs EVG TDF FTC | 0.05270 | ns |
| FTC vs TDF FTC | 0.1497 | ns |
| FTC vs Blebb | 0.01852 | ns |
| DTG TDF FTC vs EVG TDF FTC | 0.1133 | ns |
| DTG TDF FTC vs TDF FTC | 0.2103 | ** |
| DTG TDF FTC vs Blebb | 0.07909 | ns |
| EVG TDF FTC vs TDF FTC | 0.09705 | ns |
| EVG TDF FTC vs Blebb | -0.03418 | ns |
| TDF FTC vs Blebb | -0.1312 | ns |

**Table S6.**
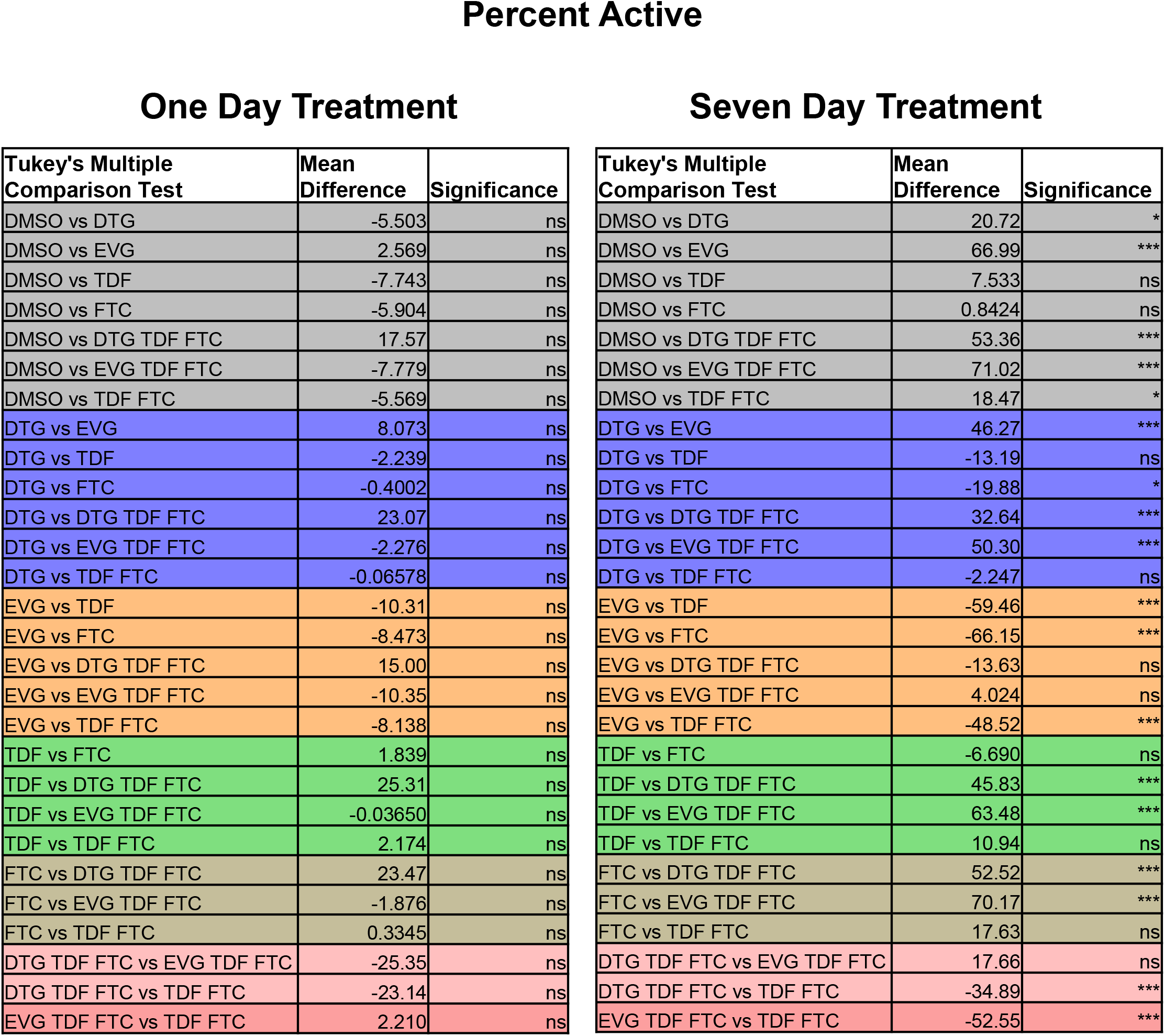
Tukey’s multiple comparison test results for percent neurons with calcium transient activity. Tables show the results of each pairwise comparison for hiPSC-neurons treated for one (left) or seven days (right) with DMSO alone, single ARVs, or combinations of ARVs (each at 10 μM). See Figure 6A, D for graph of results. ns: not significant (p> 0.05) * p < 0.05, ** p < 0.01, *** p < 0.001.

One Day Treatment
| Tukey's Multiple Comparison Test | Mean Difference | Significance |
| --- | --- | --- |
| DMSO vs DTG | -5.503 | ns |
| DMSO vs EVG | 2.569 | ns |
| DMSO vs TDF | -7.743 | ns |
| DMSO vs FTC | -5.904 | ns |
| DMSO vs DTG TDF FTC | 17.57 | ns |
| DMSO vs EVG TDF FTC | -7.779 | ns |
| DMSO vs TDF FTC | -5.569 | ns |
| DTG vs EVG | 8.073 | ns |
| DTG vs TDF | -2.239 | ns |
| DTG vs FTC | -0.4002 | ns |
| DTG vs DTG TDF FTC | 23.07 | ns |
| DTG vs EVG TDF FTC | -2.276 | ns |
| DTG vs TDF FTC | -0.06578 | ns |
| EVG vs TDF | -10.31 | ns |
| EVG vs FTC | -8.473 | ns |
| EVG vs DTG TDF FTC | 15.00 | ns |
| EVG vs EVG TDF FTC | -10.35 | ns |
| EVG vs TDF FTC | -8.138 | ns |
| TDF vs FTC | 1.839 | ns |
| TDF vs DTG TDF FTC | 25.31 | ns |
| TDF vs EVG TDF FTC | -0.03650 | ns |
| TDF vs TDF FTC | 2.174 | ns |
| FTC vs DTG TDF FTC | 23.47 | ns |
| FTC vs EVG TDF FTC | -1.876 | ns |
| FTC vs TDF FTC | 0.3345 | ns |
| DTG TDF FTC vs EVG TDF FTC | -25.35 | ns |
| DTG TDF FTC vs TDF FTC | -23.14 | ns |
| EVG TDF FTC vs TDF FTC | 2.210 | ns |

Seven Day Treatment
| Tukey's Multiple Comparison Test | Mean Difference | Significance |
| --- | --- | --- |
| DMSO vs DTG | 20.72 | * |
| DMSO vs EVG | 66.99 | *** |
| DMSO vs TDF | 7.533 | ns |
| DMSO vs FTC | 0.8424 | ns |
| DMSO vs DTG TDF FTC | 53.36 | *** |
| DMSO vs EVG TDF FTC | 71.02 | *** |
| DMSO vs TDF FTC | 18.47 | * |
| DTG vs EVG | 46.27 | *** |
| DTG vs TDF | -13.19 | ns |
| DTG vs FTC | -19.88 | * |
| DTG vs DTG TDF FTC | 32.64 | *** |
| DTG vs EVG TDF FTC | 50.30 | *** |
| DTG vs TDF FTC | -2.247 | ns |
| EVG vs TDF | -59.46 | *** |
| EVG vs FTC | -66.15 | *** |
| EVG vs DTG TDF FTC | -13.63 | ns |
| EVG vs EVG TDF FTC | 4.024 | ns |
| EVG vs TDF FTC | -48.52 | *** |
| TDF vs FTC | -6.690 | ns |
| TDF vs DTG TDF FTC | 45.83 | *** |
| TDF vs EVG TDF FTC | 63.48 | *** |
| TDF vs TDF FTC | 10.94 | ns |
| FTC vs DTG TDF FTC | 52.52 | *** |
| FTC vs EVG TDF FTC | 70.17 | *** |
| FTC vs TDF FTC | 17.63 | ns |
| DTG TDF FTC vs EVG TDF FTC | 17.66 | ns |
| DTG TDF FTC vs TDF FTC | -34.89 | *** |
| EVG TDF FTC vs TDF FTC | -52.55 | *** |

**Table S7.** Tukey’s multiple comparison test results for calcium transient event frequency. Tables show the results of each pairwise comparison for hiPSC-neurons treated for one (left) or seven days (right) with DMSO alone, single ARVs, or combinations of ARVs (each at 10 μM). See Figure 6B, E for graph of results. ns: not significant (p> 0.05) * p < 0.05, ** p < 0.01, *** p < 0.001.

One Day Treatment
| Tukey's Multiple Comparison Test | Mean Difference | Significance |
| --- | --- | --- |
| DMSO vs DTG | -0.004991 | ns |
| DMSO vs EVG | -0.02025 | ns |
| DMSO vs TDF | 0.001156 | ns |
| DMSO vs FTC | -0.01612 | ns |
| DMSO vs DTG TDF FTC | 0.005613 | ns |
| DMSO vs EVG TDF FTC | -0.01591 | ns |
| DMSO vs TDF FTC | -0.01128 | ns |
| DTG vs EVG | -0.01526 | ns |
| DTG vs TDF | 0.006147 | ns |
| DTG vs FTC | -0.01113 | ns |
| DTG vs DTG TDF FTC | 0.01060 | ns |
| DTG vs EVG TDF FTC | -0.01092 | ns |
| DTG vs TDF FTC | -0.006288 | ns |
| EVG vs TDF | 0.02141 | ns |
| EVG vs FTC | 0.004131 | ns |
| EVG vs DTG TDF FTC | 0.02587 | ns |
| EVG vs EVG TDF FTC | 0.004345 | ns |
| EVG vs TDF FTC | 0.008976 | ns |
| TDF vs FTC | -0.01728 | ns |
| TDF vs DTG TDF FTC | 0.004456 | ns |
| TDF vs EVG TDF FTC | -0.01707 | ns |
| TDF vs TDF FTC | -0.01244 | ns |
| FTC vs DTG TDF FTC | 0.02174 | ns |
| FTC vs EVG TDF FTC | 0.0002133 | ns |
| FTC vs TDF FTC | 0.004844 | ns |
| DTG TDF FTC vs EVG TDF FTC | -0.02152 | ns |
| DTG TDF FTC vs TDF FTC | -0.01689 | ns |
| EVG TDF FTC vs TDF FTC | 0.004631 | ns |

Seven Day Treatment
| Tukey's Multiple Comparison Test | Mean Difference | Significance |
| --- | --- | --- |
| DMSO vs DTG | 0.008768 | ns |
| DMSO vs EVG | 0.04941 | ** |
| DMSO vs TDF | -0.001770 | ns |
| DMSO vs FTC | -0.004350 | ns |
| DMSO vs DTG TDF FTC | 0.03159 | ns |
| DMSO vs EVG TDF FTC | 0.07615 | *** |
| DMSO vs TDF FTC | 0.003081 | ns |
| DTG vs EVG | 0.04064 | * |
| DTG vs TDF | -0.01054 | ns |
| DTG vs FTC | -0.01312 | ns |
| DTG vs DTG TDF FTC | 0.02282 | ns |
| DTG vs EVG TDF FTC | 0.06738 | *** |
| DTG vs TDF FTC | -0.005688 | ns |
| EVG vs TDF | -0.05118 | ** |
| EVG vs FTC | -0.05376 | *** |
| EVG vs DTG TDF FTC | -0.01782 | ns |
| EVG vs EVG TDF FTC | 0.02674 | ns |
| EVG vs TDF FTC | -0.04633 | ** |
| TDF vs FTC | -0.002579 | ns |
| TDF vs DTG TDF FTC | 0.03336 | ns |
| TDF vs EVG TDF FTC | 0.07792 | *** |
| TDF vs TDF FTC | 0.004851 | ns |
| FTC vs DTG TDF FTC | 0.03594 | * |
| FTC vs EVG TDF FTC | 0.08050 | *** |
| FTC vs TDF FTC | 0.007430 | ns |
| DTG TDF FTC vs EVG TDF FTC | 0.04456 | ** |
| DTG TDF FTC vs TDF FTC | -0.02851 | ns |
| EVG TDF FTC vs TDF FTC | -0.07307 | *** |

**Table S8.** Tukey’s multiple comparison test results for calcium transient mean peak amplitude. Tables show the results of each pairwise comparison for hiPSC-neurons treated for one (left) or seven days (right) with DMSO alone, single ARVs, or combinations of ARVs (each at 10 μM). See Figure 6C, F for graph of results. ns: not significant (p> 0.05) * p < 0.05, ** p < 0.01, *** p < 0.001.

One Day Treatment
| Tukey's Multiple Comparison Test | Mean Difference | Significance |
| --- | --- | --- |
| DMSO vs DTG | -1.099 | ns |
| DMSO vs EVG | 0.6888 | ns |
| DMSO vs TDF | -4.173 | ns |
| DMSO vs FTC | -1.063 | ns |
| DMSO vs DTG TDF FTC | 1.968 | ns |
| DMSO vs EVG TDF FTC | -1.797 | ns |
| DMSO vs TDF FTC | -1.659 | ns |
| DTG vs EVG | 1.787 | ns |
| DTG vs TDF | -3.074 | ns |
| DTG vs FTC | 0.03589 | ns |
| DTG vs DTG TDF FTC | 3.066 | ns |
| DTG vs EVG TDF FTC | -0.6987 | ns |
| DTG vs TDF FTC | -0.5604 | ns |
| EVG vs TDF | -4.862 | ns |
| EVG vs FTC | -1.752 | ns |
| EVG vs DTG TDF FTC | 1.279 | ns |
| EVG vs EVG TDF FTC | -2.486 | ns |
| EVG vs TDF FTC | -2.348 | ns |
| TDF vs FTC | 3.110 | ns |
| TDF vs DTG TDF FTC | 6.140 | ns |
| TDF vs EVG TDF FTC | 2.375 | ns |
| TDF vs TDF FTC | 2.514 | ns |
| FTC vs DTG TDF FTC | 3.030 | ns |
| FTC vs EVG TDF FTC | -0.7346 | ns |
| FTC vs TDF FTC | -0.5963 | ns |
| DTG TDF FTC vs EVG TDF FTC | -3.765 | ns |
| DTG TDF FTC vs TDF FTC | -3.627 | ns |
| EVG TDF FTC vs TDF FTC | 0.1383 | ns |

Seven Day Treatment
| Tukey's Multiple Comparison Test | Mean Difference | Significance |
| --- | --- | --- |
| DMSO vs DTG | 6.825 | ns |
| DMSO vs EVG | 10.96 | * |
| DMSO vs TDF | -4.487 | ns |
| DMSO vs FTC | 1.877 | ns |
| DMSO vs DTG TDF FTC | 6.521 | ns |
| DMSO vs EVG TDF FTC | 9.934 | * |
| DMSO vs TDF FTC | -2.649 | ns |
| DTG vs EVG | 4.134 | ns |
| DTG vs TDF | -11.31 | ** |
| DTG vs FTC | -4.948 | ns |
| DTG vs DTG TDF FTC | -0.3046 | ns |
| DTG vs EVG TDF FTC | 3.109 | ns |
| DTG vs TDF FTC | -9.475 | * |
| EVG vs TDF | -15.45 | *** |
| EVG vs FTC | -9.082 | ns |
| EVG vs DTG TDF FTC | -4.439 | ns |
| EVG vs EVG TDF FTC | -1.025 | ns |
| EVG vs TDF FTC | -13.61 | *** |
| TDF vs FTC | 6.364 | ns |
| TDF vs DTG TDF FTC | 11.01 | * |
| TDF vs EVG TDF FTC | 14.42 | *** |
| TDF vs TDF FTC | 1.837 | ns |
| FTC vs DTG TDF FTC | 4.643 | ns |
| FTC vs EVG TDF FTC | 8.056 | ns |
| FTC vs TDF FTC | -4.527 | ns |
| DTG TDF FTC vs EVG TDF FTC | 3.413 | ns |
| DTG TDF FTC vs TDF FTC | -9.170 | ns |
| EVG TDF FTC vs TDF FTC | -12.58 | ** |

